# Classes and continua of hippocampal CA1 inhibitory neurons revealed by single-cell transcriptomics

**DOI:** 10.1101/143354

**Authors:** Kenneth D. Harris, Hannah Hochgerner, Nathan G. Skene, Lorenza Magno, Linda Katona, Carolina Bengtsson Gonzales, Peter Somogyi, Nicoletta Kessaris, Sten Linnarsson, Jens Hjerling-Leffler

## Abstract

Understanding any brain circuit will require a categorization of its constituent neurons. In hippocampal area CA1, at least 23 classes of GABAergic neuron have been proposed to date. However, this list may be incomplete; additionally, it is unclear whether discrete classes are sufficient to describe the diversity of cortical inhibitory neurons, or whether continuous modes of variability are also required. We studied the transcriptomes of 3663 CA1 inhibitory cells, revealing 10 major GABAergic groups that divided into 49 fine-scale clusters. All previously described and several novel cell classes were identified, with three previously-described classes unexpectedly found to be identical. A division into discrete classes however was not sufficient to describe the diversity of these cells, as continuous variation also occurred between and within classes. Latent factor analysis revealed that a single continuous variable could predict the expression levels of several genes, which correlated similarly with it across multiple cell types. Analysis of the genes correlating with this variable suggested it reflects a range from metabolically highly active faster-spiking cells that proximally target pyramidal cells, to slower-spiking cells targeting distal dendrites or interneurons. These results elucidate the complexity of inhibitory neurons in one of the simplest cortical structures, and show that characterizing these cells requires continuous modes of variation as well as discrete cell classes.

---

CORTICAL CIRCUITS are composed of highly diverse neurons, and a clear definition of cortical cell types is essential for the explanation of their contribution to network activity patterns and behavior. Cortical neuronal diversity is strongest amongst GABAergic neurons. In hippocampal area CA1 – one of the architecturally simplest cortical structures – GABAergic neurons have been divided so far into at least 23 classes of distinct connectivity, firing patterns, and molecular content (Bezaire and Soltesz, 2013; Freund and Buzsaki, 1996; Klausberger and Somogyi, 2008; Pelkey et al., 2017; Somogyi, 2010; Wheeler et al., 2015). A complete categorization of CA1 inhibitory neurons would provide not only essential information to understand the computational mechanisms of the hippocampus, but also a canonical example to inform studies of more complex structures such as 6-layered isocortex.

CA1 GABAergic neurons have been divided into six major groups, based on connectivity and expression patterns of currently-used molecular markers. *Pvalb*-positive neurons (including basket, bistratified, and axo-axonic cells) target pyramidal cells’ somata, proximal dendrites or axon initial segments, firing fast spikes that lead to strong and rapid suppression of activity (Buhl et al., 1994; Hu et al., 2014). *Sst*-positive oriens/lacunosum-moleculare (O-LM) cells target pyramidal cell distal dendrites and exhibit slower firing patterns (Katona et al., 2014). GABAergic long-range projection cells send information to distal targets, and comprise many subtypes including SST*-*positive hippocamposeptal cells; NOS1-positive backprojection cells targeting dentate gyrus and CA3; and several classes of hippocamposubicular cells including trilaminar, radiatum-retrohippocampal, and PENK-positive neurons (Fuentealba et al., 2008a; Jinno, 2009; Jinno et al., 2007; Sik et al., 1994; Takács et al., 2008). CCK*-*positive interneurons are a diverse class characterized by asynchronous neurotransmitter release (Daw et al., 2009; Hefft and Jonas, 2005), that have been divided into at least 5 subtypes targeting different points along the somadendritic axis of pyramidal cells (Armstrong and Soltesz, 2012; Cope et al., 2002; Klausberger et al., 2005; Pawelzik et al., 2002; Somogyi et al., 2004). Neurogliaform and Ivy cells release GABA diffusely from dense local axons and can mediate volume transmission as well as conventional synapses (Armstrong et al., 2012; Fuentealba et al., 2008b). Interneuron-selective (I-S) interneurons comprise at least 3 subtypes specifically targeting other inhibitory neurons, and expressing one or both of *Vip* and *Calb2* (Acsady et al., 1996a, 1996b; Freund and Buzsaki, 1996; Gulyás et al., 1996). Finally, additional rare types such as large *Sst/Nos1* cells (Jinno and Kosaka, 2004) have been described at a molecular level, but their axonal targets and relationship to other subtypes is unclear.

This already complex picture likely underestimates the intricacy of CA1 inhibitory neurons. Currently defined classes likely divide into several further subtypes, and additional neuronal classes likely remain to be found (e.g. Katona et al., 2017). Furthermore, it is unclear whether a categorization into discrete classes is even sufficient to describe the diversity of cortical inhibitory neurons (Markram et al., 2004; Parra et al., 1998). For example, several *Cck* interneuron classes have been described, targeting pyramidal cells at multiple locations ranging from their somata to distal dendrites, and the molecular profile and spiking phenotype of these cells correlates with their synaptic target location, with fast-spiking cells more likely to target proximal segments of pyramidal neurons (Cope et al., 2002; Klausberger et al., 2005; Somogyi et al., 2004). Do such cells represent discrete classes with sharp inter-class boundaries, or do they represent points along a continuum? Furthermore, while a cell’s large-scale axonal and dendritic structure likely remains fixed throughout life, both gene expression and electrophysiological properties can be modified by factors such as neuronal activity (Dehorter et al., 2015; Donato et al., 2013; Mardinly et al., 2016; Spiegel et al., 2014). To what extent is the observed molecular diversity of interneurons consistent with activity-dependent modulation of gene expression?

Single-cell RNA sequencing (scRNA-seq) – which can read out the expression levels of all genes in large numbers of individual cells – provides a powerful opportunity to address these questions. This method has successfully identified the major cell classes in several brain regions (Cembrowski et al., 2016a, 2016b; Chevée et al., 2017; Ecker et al., 2017; Frazer et al., 2017; Habib et al., 2016, 2017; Macosko et al., 2015; Paul et al., 2017; Tasic et al., 2016; Usoskin et al., 2015; Zeisel et al., 2015). Nevertheless, identifying fine cortical cell classes has not been straightforward, due to both incomplete prior information on the underlying cell types, and to complicating factors such as potential continuous variability within these classes. The large body of prior work on CA1 interneurons provides a valuable opportunity to identify transcriptomic clusters with known cell types in an important cortical circuit, enabling confident identification of known and novel classes and investigation of questions such as continuous variability.

Here we describe a transcriptomic analysis of 3663 inhibitory neurons from mouse CA1. This analysis revealed 49 clusters, of which we could identify 41 with previously described cell types, with the remaining eight representing putative novel cell types. All previously described CA1 GABAergic classes could be identified in our database, but our results unexpectedly suggest that three of them are identical. The larger number of clusters occurring in our transcriptomic analysis reflected several previously unappreciated subtypes of existing classes, and tiling of continua by multiple clusters. Our data suggest a common genetic continuum exists between and within classes, from faster-firing cells targeting principal cell somata and proximal dendrites, to slower-firing cells targeting distal dendrites or interneurons. Several classes previously described as discrete represent ranges along this continuum of gene expression.

## Results

### Data collection and identification of inhibitory cells

We collected cells from six *Slc32a1-Cre;R26R-tdTomato* mice, three of age p60 and three of age p27. Cells were procured using enzymatic digestion and manual dissociation (Zeisel et al. 2015), and data were analyzed using the 10X Genomics “cellranger” pipeline, which uses unique molecular identifiers (UMIs) to produce an absolute integer quantification of each gene in each cell. The great majority of cells (4572/6971 cells total; 3283/3663 high-quality interneurons) came from the older animals. Because we observed no major difference in interneuron classes between ages, data was pooled between them (**Figure S1**). FACS sorting yielded an enriched, but not completely pure population of GABAergic neurons. A first-round clustering (using the method described below) was therefore run on the 5940 cells passing quality control, identifying 3663 GABAergic neurons (as judged by the expression of genes *Gad1* and *Slc32a1*).

### Cluster analysis

We analyzed the data using a suite of four novel algorithms, derived from a probabilistic model of RNA distributions. All four methods were based on the observation that RNA counts within a homogeneous population can be approximated using a negative binomial distribution (See methods; Lu et al., 2005; Robinson and Smyth, 2008). The negative binomial distribution accurately models the high variance of transcriptomic read counts (**Figure S2A,B**). As a consequence, algorithms based on this distribution weight the presence or absence of a gene more than its numerical expression level – for example, this distribution treats read counts of 0 and 10 as more dissimilar than read counts of 500 and 1000 (**Figure S2C**).

The algorithm we used for clustering was termed ProMMT (Probabilistic Mixture Modeling for Transcriptomics). This algorithm fits gene expression in each cluster *k* by a multivariate negative binomial distribution with cluster-specific mean **µ**_*k*_. The mean expression levels of only a small subset of genes are allowed to vary between clusters (150 for the current analysis; **Figure S3**); these genes are selected automatically by the algorithm by maximum likelihood methods. The use of such “sparse” methods is essential for probabilistic classification of high dimensional data (Bouveyron and Brunet-Saumard, 2014), and the genes selected represent those most informative for cluster differentiation. The number of clusters was chosen automatically using the Bayesian Information Criterion (BIC) (Schwarz, 1978). The ProMMT algorithm also provides a natural measure of the distinctness of each cluster, which we term the *isolation metric* (see Methods).

The ProMMT algorithm divided CA1 interneurons into 49 clusters (**Figure 1**). We named the clusters using a multilevel scheme, after genes that are strongly expressed at different hierarchical levels; for example, the cluster *Calb2.Vip.Nos1* belongs to a first level group characterized by strong expression of *Calb2* (indicating interneuron selective interneurons); a second level group *Calb2.Vip*; and a third level group distinguished from other *Calb2.Vip* cells by stronger expression of *Nos1*. This naming scheme was based on the results of hierarchical cluster analysis of cluster means, using a distance metric based on the negative binomial model (Methods; **Figure 1**).

**Figure 1.**
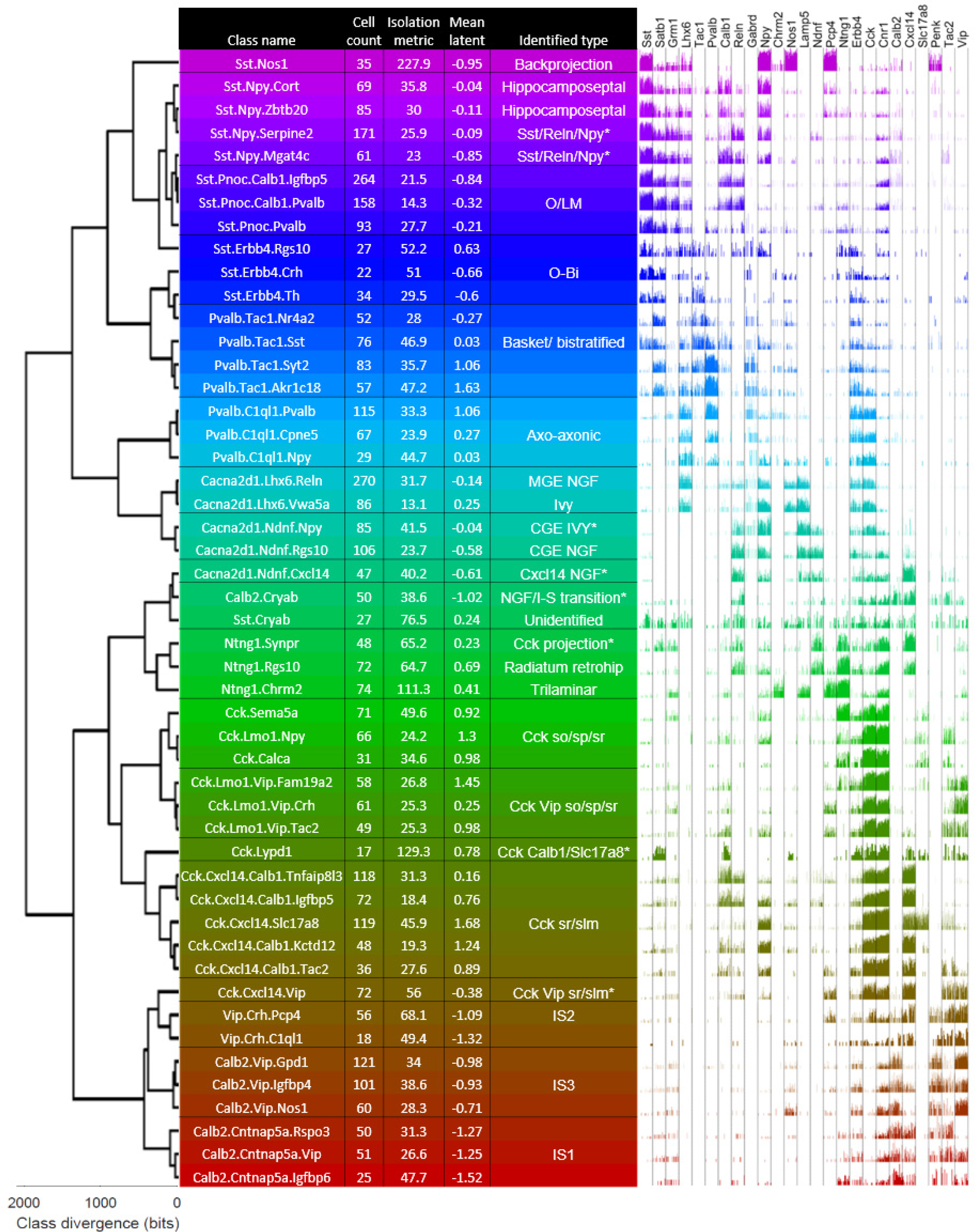
The ProMMT algorithm split CA1 GABAergic neurons into 49 clusters. Dendrogram (left) shows a hierarchical cluster analysis of these classes. Table shows class names (chosen hierarchically according to strongly expressed genes), number of cells per class, isolation metric of each class (higher for distinct classes), the mean value of latent variable analysis for cells in this class, and the biological cell type identified from its gene expression pattern. Asterisks indicating hypothesized novel classes. Right, bar chart showing log expression of 25 selected genes for all cells in the class. Note the expression pattern of *Lhx6*, which suggests a developmental origin in medial ganglionic eminence for clusters *Cacna2d1.Lhx6.Vwa5a* and above.

### Data Visualization

To visualize cell classes in two dimensions, we modified the t-stochastic neighbor embedding algorithm (Maaten and Hinton, 2008) for data with negative binomial variability, terming this approach nbtSNE. In conventional tSNE, the similarity between data points is defined by their Euclidean distance, which corresponds to conditional probabilities under a Gaussian distribution. We obtained greater separation of clusters and a closer correspondence to known cell types, by replacing the Gaussian distribution with the same negative binomial distribution used in our clustering algorithm (see Methods; **Figure S4**).

The nbtSNE maps revealed that cells were arranged in 10 major “continents” (**Figure 2**). The way expression of a single gene differed between classes could be conveniently visualized on these maps by adjusting the symbol size for each cell according to that gene’s expression level. Consistent with previous transcriptomic analyses, we found that classes were rarely if ever identified by single genes, but rather by combinatorial expression patterns. Thanks to the extensive literature on CA1 interneurons, 25 genes together sufficed to identify the main continents with known cell classes (**Figure 3**), and it was also possible to identify nearly all the finer subclasses using additional genes specific to each class (**Supplementary Text**).

**Figure 2.**
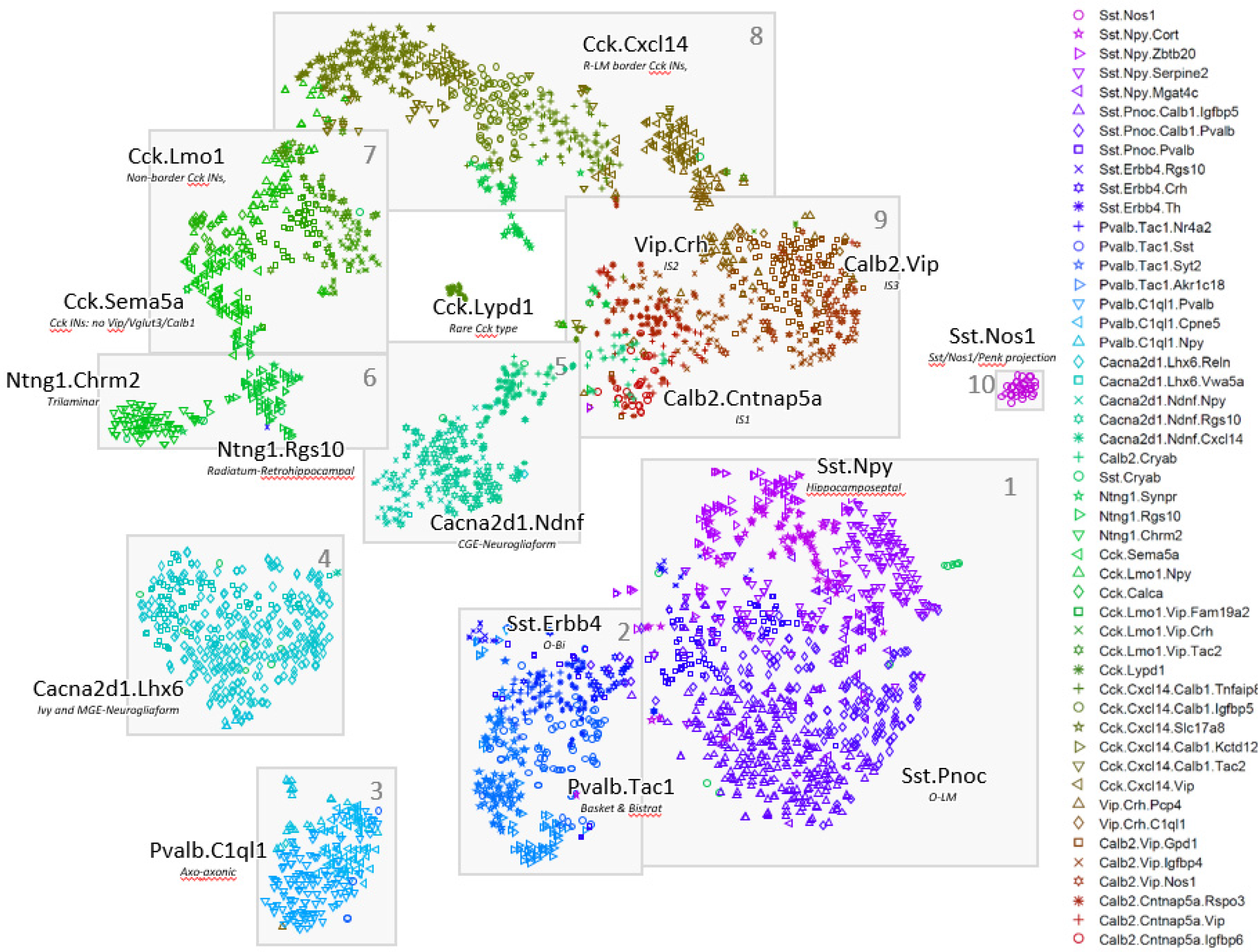
Two-dimensional visualization of expression patterns using nbtSNE algorithm, which places cells of similar expression close together. Each symbol represents a cell, with different color/glyph combinations representing different cell classes (legend, right). Grey boxes and numbers refer to the “continents” referred to in the text and subsequent figures.

**Figure 3.**
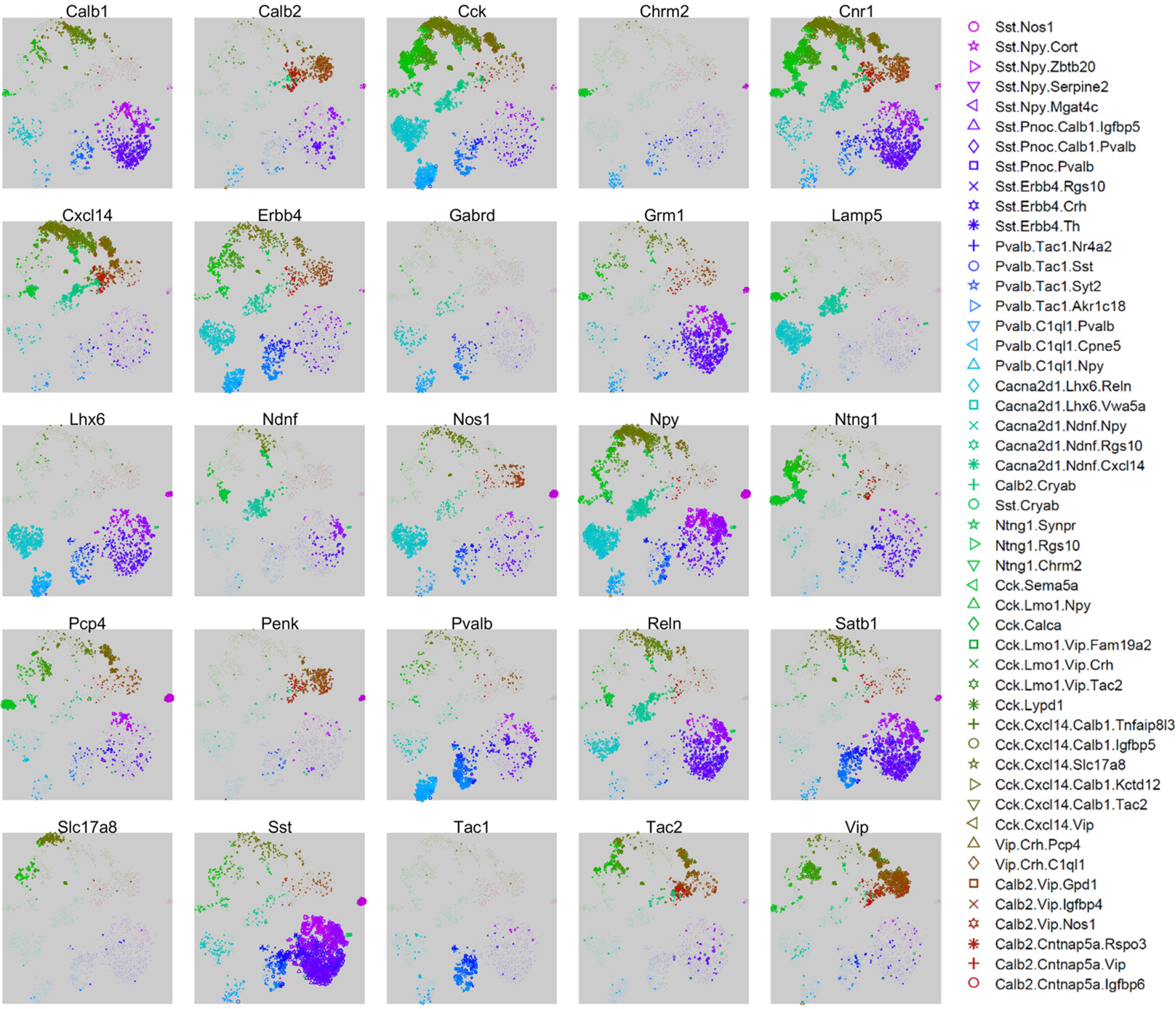
Expression levels of 25 selected genes, that together allow identification of major cell classes. Each subplot shows an nbtSNE map of all cells, with marker size indicating log-expression level of the gene named above the plot.

### Identification of cell types

Previous work has extensively characterized the connectivity, physiology, and firing patterns of CA1 inhibitory neurons, and these cellular properties have been related to expression of large numbers of marker genes. We next sought to identify our transcriptomic clusters with previously defined cell types, taking advantage of the “Rosetta Stone” provided by this extensive prior research. Explaining how the identifications were made requires an extensive discussion of the previous literature, which is presented in full in the Supplementary Text. Here, we briefly summarize the major subtypes identified (summarized in **Figure 4)**.

**Figure 4:**
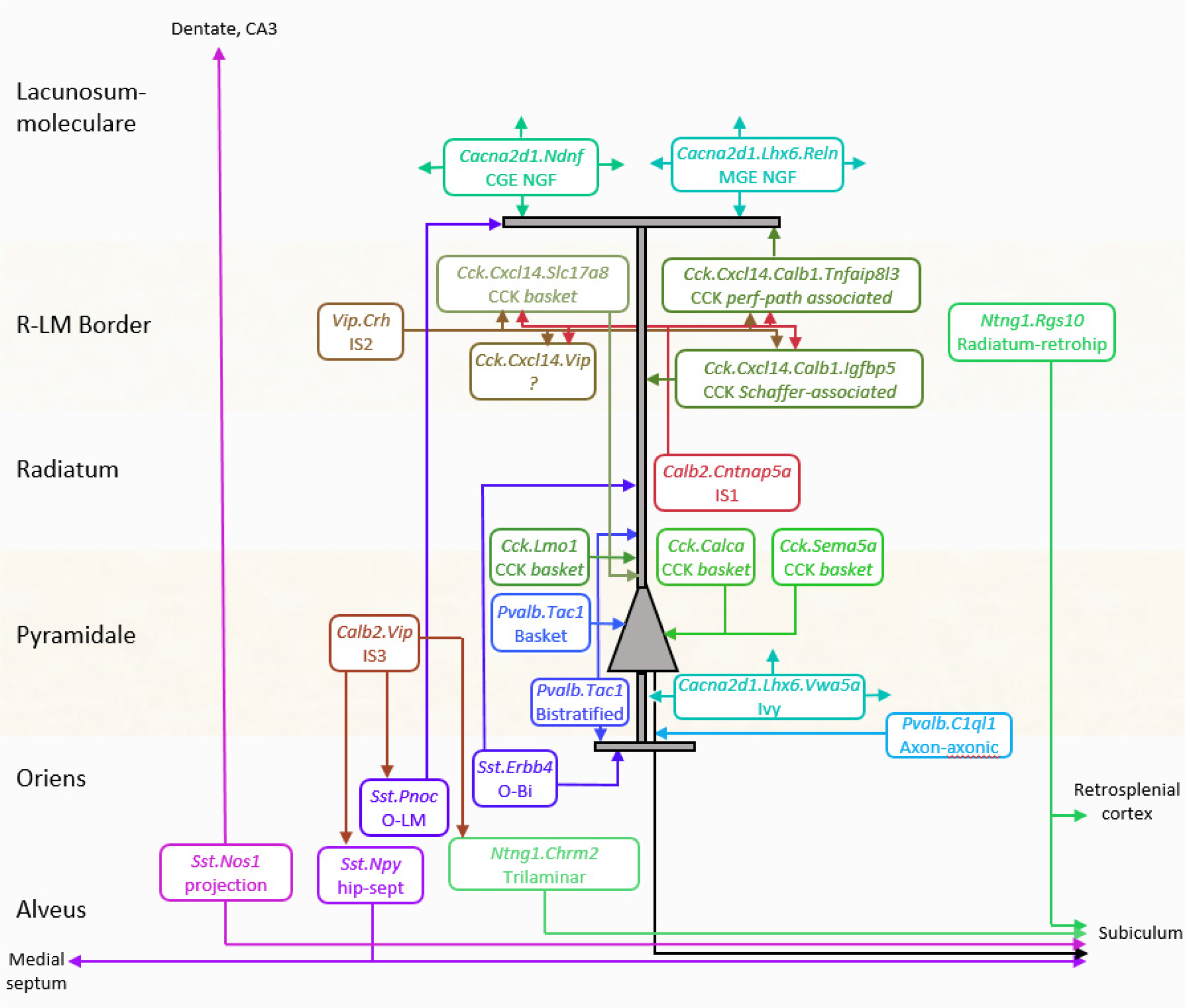
Inferred circuit diagram of identified GABAergic cell types. The identification of transcriptomic clusters with known cell classes is described in full in the supplementary material. Laminar locations and connections between each class are derived from previous literature.

Continent 1 was identified with the *Sst* positive hippocamposeptal and O-LM cells of *stratum oriens (so)*. These cells all expressed *Sst* and *Grm1*, and were further divided into two *Npy+/Ngf+* clusters identified as hippocamposeptal neurons (Acsády et al., 2000), and three *Pnoc+/Reln+/Npy-*clusters identified with O-LM cells (Katona et al., 2014). In addition, Continent 1 contains a previously undescribed subclass positive for *Sst, Npy*, and *Reln*.

Continent 2 was identified as basket and bistratified cells. These were all positive for *Tac1* (the precursor to the neuropeptide Substance P), as well as *Satb1* and *Erbb4*, but were negative for *Grm1*. They were divided into two *Pvalb+/Sst-*clusters identified with basket cells, two *Pvalb+/Sst+/Npy+* clusters identified with bistratified cells (Klausberger et al., 2004), and three *Pvalb-*clusters identified with Oriens-Bistratified (o-Bi) cells (Losonczy et al., 2002).

Continent 3 was identified as axo-axonic cells, due to their expression of *Pvalb* but not *Satb1* (Viney et al., 2013) This continent’s three clusters were *Tac1-*negative but positive for other markers including *Snca, Pthlh* and *C1ql1*, which have also been associated with axo-axonic cells in isocortex (Paul et al., 2017; Tasic et al., 2016). We note that this dichotomy of *Pvalb* interneurons into *Tac1* positive and negative subclasses is likely homologous to previous observations in isocortex (Vruwink et al., 2001).

Continent 4 was identified as Ivy cells and MGE-derived neurogliaform cells. These cells expressed *Cacna2d1*, which we propose as a unique identifier of hippocampal neurogliaform/ivy cells, as well as *Lhx6* and *Nos1* (Tricoire et al., 2010). They were divided into a *Reln+* cluster identified with MGE-derived neurogliaform cells, and a *Reln-/Vwa5a+* cluster identified with Ivy cells (Fuentealba et al., 2008b). This continent is homologous to the isocortical *Igtp* class defined by Tasic et al (2016), which we hypothesize may represent isocortical neurogliaform cells of MGE origin; this hypothesis could be confirmed using fate-mapping.

Continent 5 was identified as CGE-derived neurogliaform cells. Its three clusters contained *Cacna2d1* and many other genes in common with those of continent 4, but lacked *Lhx6* and *Nos1* (Tricoire et al., 2010). Similar to isocortical putative neurogliaform cells, this continent expressed *Ndnf* and contained a distinct subtype positive for *Cxcl14* (Tasic et al., 2016). As with continent 4, continent 5 mainly expressed *Reln* but also contained a small *Reln-*negative cluster, which we suggest forms a rare and novel class of CGE-derived ivy cell.

Continent 6 was identified with *Sst-*negative long-range projection interneurons. It divided into two distinct clusters, both of which were strongly positive for *Ntng1*. The first strongly expressed *Chrm2* but lacked *Sst* and *Pvalb*, identifying them as trilaminar cells (Ferraguti et al., 2005; Jinno et al., 2007). The second subgroup lacked most classical molecular markers; this fact, together with their inferred laminar location at the *sr/slm* border, identified them as putative radiatum-retrohippocampal neurons that project to retrosplenial cortex (Jinno et al., 2007; Miyashita and Rockland, 2007).

Continents 7 and 8 were identified as what are traditionally called *Cck* interneurons. This term is somewhat unfortunate: while these cells indeed strongly express *Cck*, many other inhibitory classes express *Cck* at lower levels, including even *Pvalb+* basket cells (Tricoire et al., 2011). Continents 7 and 8 cells comprised thirteen highly diverse clusters, but shared strong expression of *Cnr1, Sncg, Trp53i11* and several other novel genes. Continent 8 is distinguished by expression of *Cxcl14*, which localizes these cells to the border of *stratum radiatum* and *stratum lacunosum-moleculare (sr/slm)*. This continent comprised a continuum ranging from soma-targeting basket cells identified by their *Slc17a8* (vGlut3) expression, to dendrite targeting cells identified by expression of *Calb1* or *Reln* (Klausberger et al., 2005; Somogyi et al., 2004). Continent 7, lacking *Cxcl14*, was identified as *Cck* cells of other layers, and contained multiple subtypes characterized by the familiar markers *Calb1, Vip*, *Slc17a8* (Somogyi et al., 2004), as well novel markers such as *Sema5a* and *Calca*. Associated with continent 8 were several apparently novel subtypes: a rare and distinct group positive for both *Scl17a8* and *Calb1* and marked nearly exclusively by *Lypd1*; a *Ntng1+/Ndnf+* subgroup related to cells of continent 6; and a group strongly expressing both *Vip* and *Cxcl14*, which therefore likely corresponds to a novel *Vip+/Cck+* interneuron at the *sr/slm* border.

Continent 9 was identified as interneuron-selective interneurons. Its eight clusters fell into three groups: *Calb2+/Vip-*neurons identified as IS-1 cells; *Calb2-/Vip+* neurons identified as IS-2 cells; and *Calb2+/Vip+* neurons identified as IS-3 cells (Acsady et al., 1996a; Freund and Buzsaki, 1996; Gulyás et al., 1996; Tyan et al., 2014). All expressed *Penk* (Blasco-Ibanez et al., 1998). These cells contained at least two novel subgroups: an IS-3 subtype positive for *Nos1* and *Myl1*, homologous to the *Vip Mybpc2* class defined in isocortex (Tasic et al., 2016); and a rare subclass of IS-1 cell positive for *Igfbp6*.

Continent 10 contained a single highly distinct cluster located in an “island” off continent 1. In contained cells strongly positive for *Sst* and *Nos1* (Jinno and Kosaka, 2004), whose expression pattern is consistent with that of both backprojection cells (Sik et al., 1994) and PENK-positive projection cells (Fuentealba et al., 2008a), suggesting that these three previously-identified classes reflect a single cell type.

### Comparison with isocortical classes

Our finding of 49 clusters in a sample of 3663 CA1 cells contrasts with a previous study of isocortical area V1, which found 23 clusters from a sample of 761 inhibitory neurons (Tasic et al., 2016). One can imagine three reasons for the greater number of clusters found in the present study: the larger sample size used here may have resulted in our resolving more clusters; the use of a different clustering algorithm may have allowed the current study to reveal finer cell types; or, area CA1 might genuinely contain more diverse inhibitory neurons than isocortex. To address these questions, we performed two analyses. First, we applied our clustering algorithm to the data of Tasic et al (2016); and we re-analyzed subsamples of the data of both the current study and of Tasic et al (2016) to see how the number of clusters found varies with cell count and with sequencing depth.

Applying the ProMMT algorithm to the Tasic dataset yielded 30 clusters (**Supplementary figure S5A,B**). The cluster assignments almost completely overlapped as far as top-level groupings, but showed some more subtle distinctions in finer levels clusters (**Supplementary figure S5C**). We examined 3 of these novel classes differences in more depth, to ask whether the finer distinctions found by the ProMMT algorithm could correspond to genuine biological cell classes. The most notable of these was cluster 11, which contained neurons that had previously been assigned to the neurogliaform clusters Ndnf Cxcl14, Ndnf Car4, but lacked common neurogliaform markers such as *Lamp5* and *Gabrd*. Instead, cells in these clusters expressed *Calb2* and *Penk* but not *Vip*, suggesting interneuron-selective cells homologous to hippocampal IS-1 cells, and potentially matching the *Vip-*negative interneuron-selective layer 1 “single-bouquet cells” (SBCs) described Jiang et al (Jiang et al., 2013, 2015). To test whether cluster 11 indeed corresponds to SBCs, we took advantage of a Patch-seq study (Cadwell et al., 2016) that contrasted gene expression in anatomically identified layer 1 SBCs and neurogliaform cells (**Supplementary Figure S5D**). We found that the genes that Cadwell et al had reported as distinguishing SBCs from neurogliaform cells indeed occurred in almost entirely non-overlapping populations of cells; furthermore, these populations closely matched the ProMMT clusters identified with SBCs and neurogliaform cells. Examination of two further subdivisions found by the ProMMT algorithm again revealed genes uniquely expressed in non-overlapping subpopulations of the Sst Cbln4 and Vip Parm1 clusters (**Supplementary Figure S6**). We conclude that the larger number of clusters identified by the ProMMT algorithm at least in part results from its ability to distinguish subtle variations in gene expression between related cell types.

To ask whether the greater number of clusters found in the current study might in part arise from its larger sample size, we reran the cluster analysis on randomly-selected subsets of cells from our dataset. We found a strong linear increase in the number of clusters found with sample size (**Supplementary figure S7A**). To investigate what effect sequencing depth may have had, we resampled our dataset to simulate lower read counts for the same cells, and again found an approximately linear increase in the number of identified clusters with read count (**Supplementary figure S7B,C**). We performed similar analyses on Tasic et al’s data, and obtained similar results (**Supplementary Figure S7D,E**).

We therefore conclude that the larger number of clusters found by the current study is more likely to reflect a combination of larger sample size and more sensitive clustering algorithms, than a greater number of biological cell types in CA1 than in V1. Furthermore, we expect that an even larger sample size or greater sequencing depth would have revealed yet more, finely distinguished cell types.

### Continuous variation between and within cell classes

Although the major continents of the expression map were clearly separated, clusters within these continents often appeared to blend into each other continuously. This suggests continuous gradation in gene expression patterns: while our probabilistic mixture model will group cells from a common negative binomial distribution into a single cluster, it will tile cells of continuously graded mean expression into multiple clusters.

Although visualization methods such as nbtSNE can suggest whether classes are discrete or continuously separated, they are not sufficient to confirm the suggestion. Such methods exhibit local optima, raising the possibility that apparent continuity only occurs for particular initialization conditions. Furthermore, as nbtSNE is based on a subset of genes, it is conceivable that discrete/continuous patterns occur only for this subset.

To confirm the apparent continuity or discreteness of these groups, we therefore employed a novel method of negative binomial discriminant analysis, that is independent of nbtSNE and considers all genes. Given a pair of cell classes, this method compares how close each cell’s whole-genome expression pattern is to each class, using a cross-validated likelihood ratio statistic. For two classes identified as basket and axoaxonic cells, the histogram of likelihood ratios was clearly bimodal (**Figure 5A**, top), indicating that every cell exhibited a much stronger fit to its own class than to the other, and confirming the discrete separation of these classes. A second example of clusters identified with Ivy and MGE-neurogliaform cells however showed different behavior (**Figure 5A**, middle): a unimodal likelihood ratio histogram indicated that the two clusters ran smoothly into each other, tiling a continuum of gene expression patterns. The bimodality of the likelihood ratio can be captured by a d’ statistic, which for these two examples was 7.2 and 1.5, respectively. Perhaps ironically, the degree to which two neighboring classes are discrete or continuous was itself a continuous variable. For example, *Slc17a8-*expressing *Cxcl14/Cck* neurons showed largely continuous overlap with their neighboring *Cck/Cxcl14* cells, but with some small indication of bimodality, characterized by a d’ of 3.1 (**Figure 5A**, bottom). We conclude that while truly discrete cluster separations do exist, the dataset is not fully described as a set of discrete classes, and that many clusters tile continuous dimensions (**Figure 5B**).

**Figure 5.**
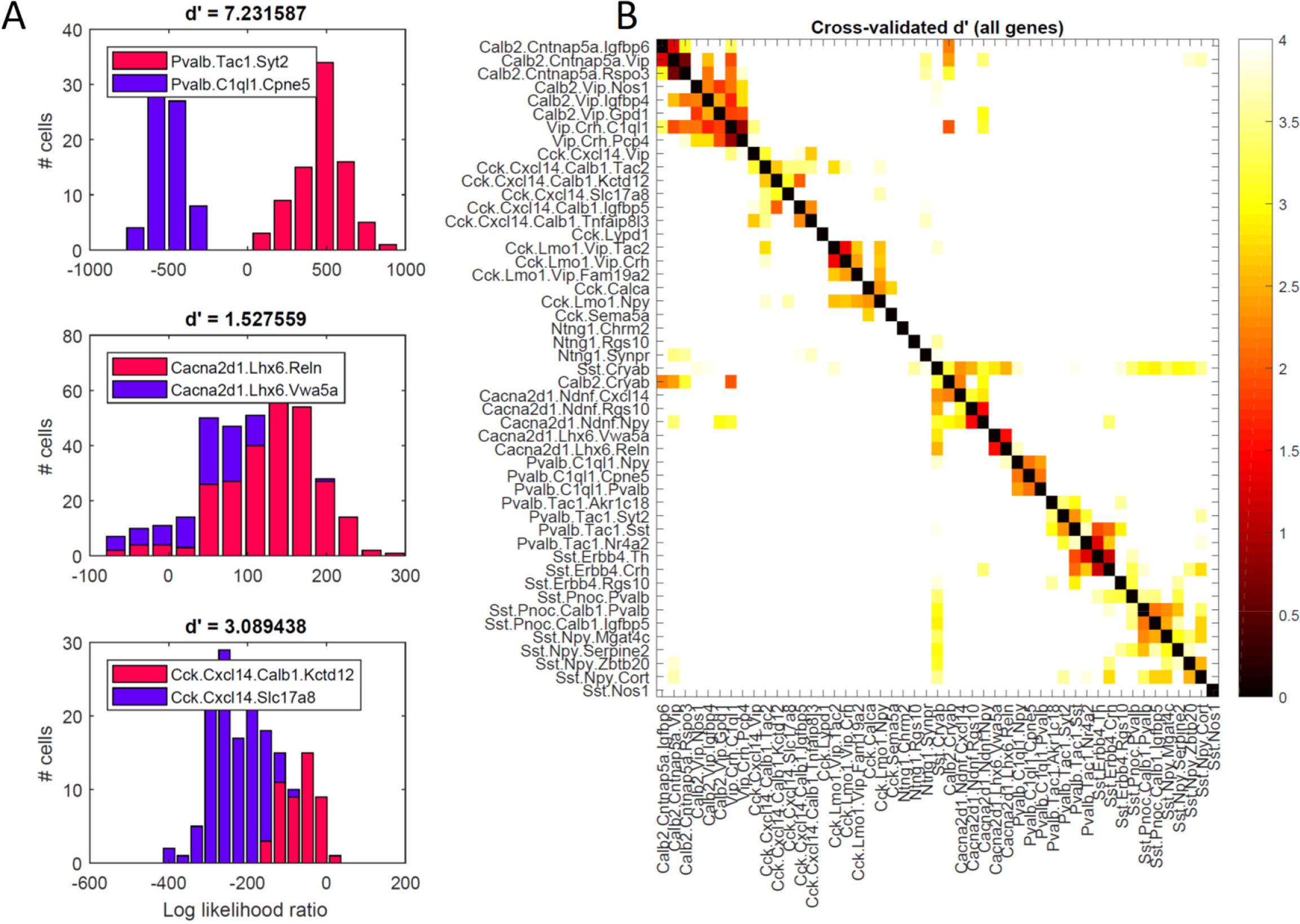
Analysis of discrete vs. continuous variation by negative binomial discriminant analysis. **A**, Histogram of log-likelihood ratios for three example cluster pairs, measuring how much better each cell’s whole-genome expression pattern is explained by one or the other clusters. The top histogram (basket vs. axo-axonic cells) is clearly bimodal, indicating clearly discrete separation. The bottom two histograms (ivy vs. MGE-neurogliaform cells; two subclasses of Cck/Cxcl14 cells) show substantial overlap, indicating continuous variation between clusters. The degree of bimodality is captured by the d’ statistic, above each plot. **B**, Pseudocolor matrix showing continuity of each pair of clusters, as assessed by d’ statistic. White means strongly bimodal, darker colors indicate continuity.

### Latent factor analysis reveals a common mode of variation across all cell types that correlates with axon target location

The existence of continuous variation in gene expression suggests that cluster analysis is not giving a complete picture of neuronal gene expression patterns. To further study the biological significance of continuously varying gene expression, we therefore applied a complementary method, latent factor analysis. Cluster analysis can be viewed as an attempt to summarize the expression of all genes using only a single discrete label per cell (the cell’s cluster identify), where the value this label takes for each cell is not directly observed, but “latent” and inferred from the data. Latent factor analysis also attempts to predict the expression of all genes using only a single variable (the “latent factor”), but now with a continuous rather than discrete distribution. As with cluster analysis, the latent factor is not directly observed, but is inferred for each cell. Latent factor analysis operates without knowledge of cluster identity, and therefore requires that the same rules be used to predict gene expression from the latent factor, for cells of all types. Clearly, one should expect neither method to precisely predict the expression of all genes from a single variable; but the rules of cellular organization they reveal may provide important biological information.

As expected, latent factor analysis produced a complementary view to cluster analysis (**Figure 6A**). Knowing a cell’s cluster identity did not suffice to predict its latent factor value, and *vice versa*. For example, the ranges of latent factor values for cells in the clusters identified with *Cck* and *Pvalb* basket cells overlapped. Nevertheless, the range of possible latent factor values was not identical between clusters, and the mean latent factor value of each cluster differed in a manner that had a clear biological interpretation.

**Figure 6.**
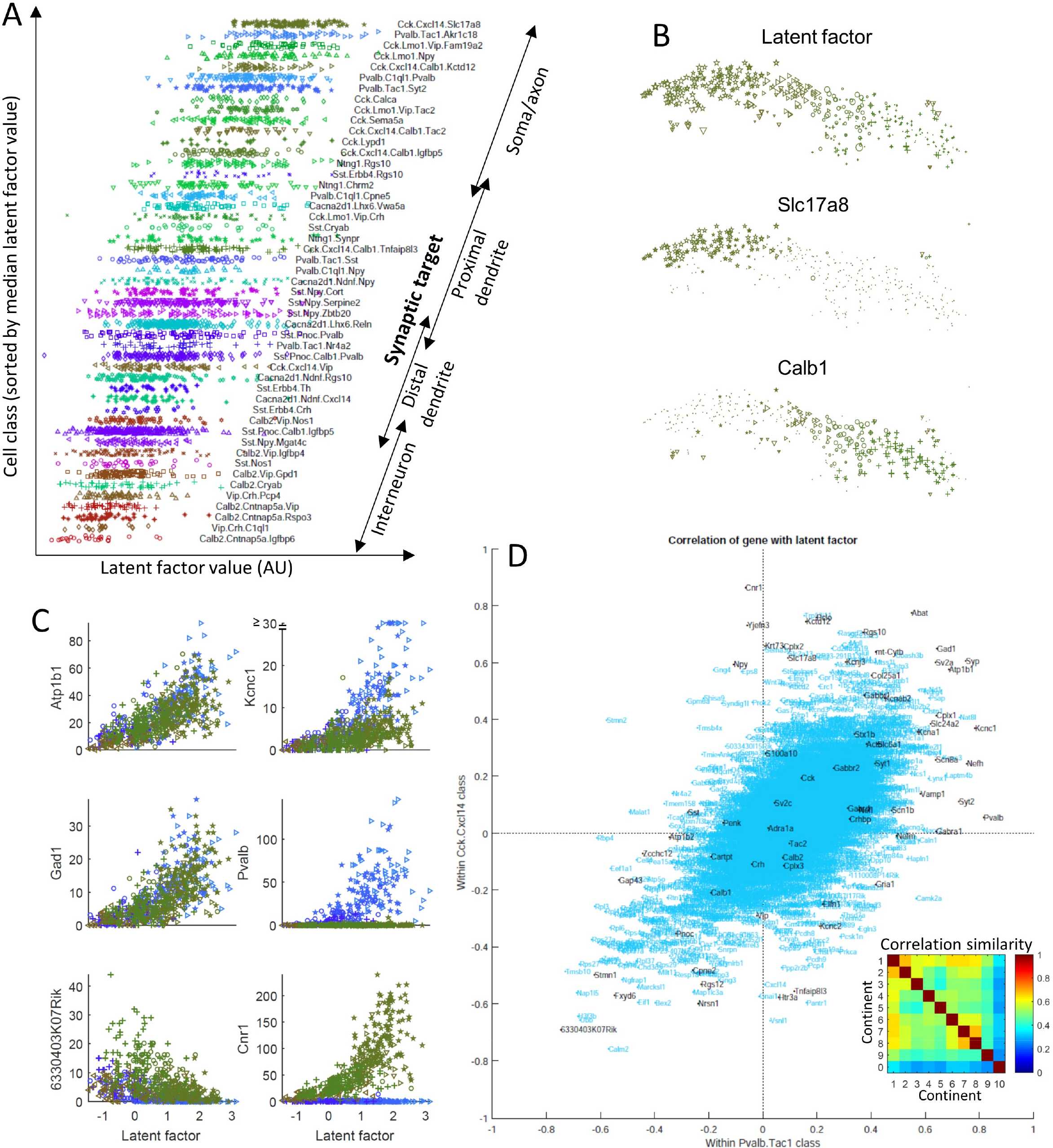
Latent factor analysis reveals a common mode of continuous variation that is consistent across cell types and correlates with axon target location. **A**, Latent factor analysis assigns a single number to each cell, via a search for the factor values that best predict expression of multiple genes. Mean factor values within each cluster differ systematically in a way that correlates with the identified cell class’ axon target location. Each point represents a cell, with x-coordinate showing latent factor value and y-coordinate showing cluster, sorted by mean latent factor value. **B**, Continuous gradient of Latent factor values across continent 8 (top; symbol size denotes latent factor value). Largest values are found in western *Slc17a8-*expressing neurons identified with soma-targeting *Cck* basket cells; smallest values found in eastern *Calb1-*expressing neurons identified with dendrite targeting *Cck* cells **C**, Correlation of latent factor values with expression of 6 example genes. Symbols as in (A); blue, continent 2 (basket/bistratified), green, continent 8 (*Cck/Cxcl14)*. **D**, Correlations of genes with the latent factor are preserved across cell classes. X-axis: correlation of gene with latent factor in continent 2; y-axis, correlation in continent 8; Spearman’s *ρ*=0.58, p<10^−100^. Inset, Spearman *ρ* values for all pairs of continents, p<10^−100^ in each case.

The mean latent factor value of each cluster correlated with the axon target location of the corresponding cell type (**Figure 6A**). The clusters showing largest mean latent factor values were identified with soma-targeting basket cells (both *Pvalb* and *Cck* expressing), and with axo-axonic cells. Lower values of the latent factor were found in clusters identified with dendrite-targeting *Cck* cells, and with bistratified, Ivy and hippocamposeptal cells, which target pyramidal cells’ proximal dendrites (Fuentealba et al., 2008b; Takács et al., 2008). Still lower values of the latent factor were found in clusters identified with neurogliaform and O-LM cells, which target pyramidal distal dendrites. The lowest values of all were found in clusters identified with cells synapsing on inhibitory interneurons: the IS cells of continent 9, and the Sst/Penk/Nos1 cells of continent 10, whose local targets are *Pvalb* cells (Fuentealba et al., 2008a).

While mean values of the latent factor differed between continents, there was also substantial variability within cells of a single continent. For example, a gradient of latent factor values was seen within continent 8 (identified with *Cck-*positive neurons at the *sr/slm* border), with larger values in the west smoothly transitioning to smaller values in the east (**Figure 6B**). Comparison of gene expression patterns in continent 8 to previous work again suggested that this gradient in latent factor values correlates with axon target location. Indeed, immunohistochemistry has demonstrated that CCK*-*positive cells expressing SLC17A8 (expressed in western continent 8) project to the pyramidal layer (Somogyi et al., 2004), while those expressing CALB1 (expressed in the east) target pyramidal cell dendrites (Cope et al., 2002; Gulyás and Freund, 1996; Klausberger and Somogyi, 2008). The cannabinoid receptor *Cnr1*, which is more strongly expressed in soma-targeting neurons (Dudok et al., 2015; Lee et al., 2010) was also more strongly expressed in western cells with larger latent factor values.

As expected, the expression levels of many individual genes correlated with the latent factor, furthermore the directions of these correlations were consistent even within distantly related cell types. We investigated the relationships of genes to the latent factor by focusing initially on the *Pvalb* and *Cck* expressing cells of continents 2 and 8 (**Figure 6C**). Most genes correlated similarly with the latent factor in both classes. For example, the Na^+^/K^+^ pump *Atp1b1* and the GABA synthesis enzyme *Gad1* correlated positively with the latent factor for multiple cell types, while *6330403K07Rik*, a gene of unknown function, correlated negatively. Some genes’ expression levels depended on both cell type and latent factor value. For example, the ion channel *Kcnc1* (which enables rapid action potential repolarization in fast-spiking cells) correlated positively with the latent factor in both *Pvalb* and *Cck* cells, but its expression was stronger in *Pvalb* cells, even for the same latent factor value. Other genes showed correlations with the latent factor, but only within the specific classes that expressed them. For example, expression of *Pvalb* correlated with the latent factor within cells of continent 2 but the gene was essentially absent from cells of continent 8; conversely, *Cnr1* expression correlated with the latent factor in continent 8, but was essentially absent in cells of continent 2. Thus, the latent factor value is not alone sufficient to predict a cell’s gene expression pattern, but provides a summary of continuous gradation in the expression of multiple genes in multiple cell types.

The relationship of genes to latent factor values was statistically similar across cell types. To demonstrate this, we computed the Spearman correlation of each gene’s expression level with the latent factor, separately within cells of each continent (Supplementary Table 1). As expected from the scatterplots (**Figure 6C**), the correlation coefficients for *Atp1b1, Gad1*, and *6330403K07Rik* were similar between continents 2 and 8 (Figure 6D). Also as expected, *Pvalb* and *Cnr1* showed strong positive correlations with the latent factor within the continent where these genes were expressed, but correlations close to zero within the continent where they were barely expressed. In general, the correlation coefficients of genes with the latent factor were preserved between continents 2 and 8 (**Figure 6D;** Spearman rank correlation ρ= 0.58, p<10^−100^). A similar relationship was found across all continents (Figure 6D, inset; p<10^−100^ in each case), although cells of continents 9 and 10 showed less similarity than continents 1-8. Furthermore, similar results were obtained when analyzing isocortical data, most notably in isocortical *Pvalb* cells (**Supplementary Figure S8**)

In summary, the expression of many genes correlates with a single continuous variable, the latent factor value assigned to each cell. While this latent factor does not provide a complete summary of a cell’s gene expression pattern, the direction and strength of the correlation of individual genes to the latent factor is largely preserved across cell types. Furthermore, while a cell’s latent factor value was not simply a function of its cell class, mean latent factor values differed between clusters, being largest for clusters identified with cell types whose axons target pyramidal somata or axon initial segments, and smallest for clusters identified with cell types targeting pyramidal distal dendrites or interneurons.

### Biological significance of genes correlating with the latent factor

The above results suggest that the expression of a large set of genes is modulated in a largely consistent way across multiple cell types, in a manner that correlated with their axonal targets. What biological functions might these genes serve? While one might certainly expect structural genes be differentially expressed between soma- and dendrite-targeting interneurons, these cells also differ in their physiology. Indeed, *Pvalb*-expressing basket cells are known for their fast spiking phenotype, which produces rapid, powerful perisomatic inhibition, and is mediated by a set of rapidly-acting ion channels and synaptic proteins including *Kcnc1, Kcna1, Scn1a, Scn8a*, and *Syt2* (Hu et al., 2014). Although most other interneurons show regular-spiking phenotype, CCK-expressing basket cells with a fast-spiking phenotype have also been reported (Cope et al., 2002; Pawelzik et al., 2002). We therefore hypothesized that genes responsible for the fast-spiking phenotype might be positively correlated with the latent factor, due to increased expression in soma-targeting cells of all classes.

Consistent with this hypothesis, genes associated with fast-spiking phenotype (*Kcnc1, Kcna1, Scn1a, Scn8a, Syt2)* were amongst the genes most positively correlated with the latent factor in both *Pvalb* and *Cck* basket cells (**Figure 6D**). However, this positive correlation was not restricted to these cell types: in an ordering of the correlations of all genes with the latent factor (taking into account cells of all types), these genes ranked in the 99.9^th^, 98.3^rd^, 99.5^th^, 98.9^th^, and 95^th^ percentiles respectively (**Supplementary table 1**).

Other gene families positively correlated with the latent factor included genes associated with mitochondria (e.g. *mt-Cytb*), ion exchange and metabolism (e.g. *Atp1b1; Slc24a2*), GABA synthesis and transport (e.g. *Gad1, Slc6a1*), vesicular release (e.g. *Syp, Sv2a, Cplx2, Vamp1*), fast ionotropic glutatmate and GABA receptors (e.g. *Gria1, Gabra1*), as well as GABAB receptors (e.g. *Gabbr1, Gabbr2, Kcnj3, Kctd12*). The genes correlating negatively with the latent factor were less familiar, but included *Atp1b2*, a second isoform of the Na^+^/K^+^ pump; *Fxyd6*, which modulates its activity; *Nrsn1*, whose translation is suppressed after learning (Cho et al., 2015), as well as many neuropeptides (*e.g. Sst, Vip, Cartpt, Tac2, Penk, Crh;* exceptional neuropeptides such as *Cck* showed positive correlation). Genes associated with neurofilaments and intermediate filaments (e.g. *Nefh, Nefl, Krt73*) tended to show positive weights, while genes associated with actin processing (e.g. *Gap43, Stmn1, Tmsb10*) tended to show negative weights. Many other genes of as yet unknown function correlated positively and negatively with the latent factor (for example *6330403K07Rik*). Relating the latent factor correlations of each gene to their Gene Ontology annotations (which are not granular enough to list annotations such as fast-spiking physiology), suggested that negatively correlated genes tended to be associated with translation and ribosomes, while positively correlated genes were associated with diverse functions including transcription, signal transduction, ion transport, and vesicular function and associated with cellular compartments including mitochondria, axons, and dendrites (**Supplementary table 2**).

We therefore suggest that cells with large values of the latent factor not only target more proximal components of pyramidal cells, but also express genes enabling a faster spiking firing pattern, more synaptic vesicles and larger amounts of GABA release; receipt of stronger excitatory and inhibitory inputs; and faster metabolism. These are all characteristics of *Pvalb-*expressing fast-spiking interneurons (Hu et al., 2014), but a similar continuum was observed within all cell types, suggesting that these genes are commonly regulated in all CA1 interneurons.

The fact that the latent factor differs systematically between cells with different axonal targets suggests that this property is in good measure fixed, as it seems unlikely that neurons would make major changes to their axonal targets in adulthood. Nevertheless, interneuronal gene expression can be modulated by activity, and some of the genes that were most strongly correlated with the latent factor (*Pvalb, Kcna1*) are amongst those with activity-dependent modulation (Cohen et al., 2016; Dehorter et al., 2015; Donato et al., 2013; Mardinly et al., 2016; Spiegel et al., 2014).

To investigate whether the genes correlated with the latent factor might also be partially modulated by neuronal activity, we correlated each gene’s latent factor score with that gene’s modulation by *in vivo* light exposure after dark housing, using data from three classes of visual cortical interneurons (made available by Mardinly et al., 2016). We observed a moderate relationship of latent factor weighting to activity modulation in *Sst* neurons (r=.26; p<10^−12^; **Supplementary Figure S9**), suggesting that activity dependent modulation of *Sst* cells may cause them to move along the continuum of latent factor values. A weaker but still significant correlation was observed for *Pvalb* neurons (r=0.11; p<0.002), whereas no significant relationship was found for *Vip* neurons (p=0.17). These data therefore suggest that a portion of the continuous variability of gene expression observed in CA1 interneurons may arise from activity-dependent modulation, but that such modulation is unlikely to be a full explanation for the genetic continua revealed by latent factor analysis.

### Histological confirmation of transcriptomic predictions

The transcriptomic classification we derived makes a large number of predictions for the combinatorial expression patterns of familiar and novel molecular markers in distinct CA1 interneuron types. To verify our transcriptomic classification, we set out to test some of these predictions using traditional methods of molecular histology.

Our first tests regarded the very distinct *Sst.Nos1* cluster of continent 10. This cluster’s expression pattern matched three previously reported rare hippocampal inhibitory cell types: large SST-immunopositive cells that are intensely immunoreactive for NOS1 throughout the cytoplasm revealing their full dendrites (Jinno and Kosaka, 2004); PENK-positive projection cells (Fuentealba et al., 2008a); and strongly NADPH diaphorase-labelled (i.e. NOS1 positive) backprojection cells (Sik et al., 1994). We therefore hypothesized that these cell types, previously regarded as separate, may in fact be identical. To test this hypothesis, we performed a series of triple and quadruple immunoreactions, focusing on the intensely NOS1-positive neurons (n=3 mice, n=70 cells: 39% in *so/*alveus; 10% in *sp*; 27% in *sr*; 24% at *sr/slm* border). Similar to previously reported PENK-projection, backprojection, and SST/NOS1 cells (Fuentealba et al., 2008a; Jinno and Kosaka, 2004; Sik et al., 1994) – but unlike SST-positive O-LM cells (Katona et al., 2014) – these neurons all showed aspiny or sparsely spiny dendrites. As expected from the *Sst.Nos1* cluster, we found that they were all SST/NPY double positive (n=20/20) and were virtually all weakly positive for CHRM2 (n=36/38) and GRM1 (n=17/17) in the somato-dendritic plasma membrane, strongly positive for PCP4 (n=19/21) in the cytoplasm and nucleus, and for PENK (n=35/42) in the Golgi apparatus and granules in the soma and proximal dendrites (**Figure 7**). By contrast, the more numerous moderately NOS1 positive cells (which include many interneuron types such as ivy, MGE-neurogliaform and a subset of IS-3 neurons) were mostly immunonegative for CHRM2, PCP4 and PENK, although some were positive for GRM1. Our results are therefore consistent with the hypothesis that all three previously reported classes correspond to the *Sst1.Nos1* cluster.

**Figure 7.**
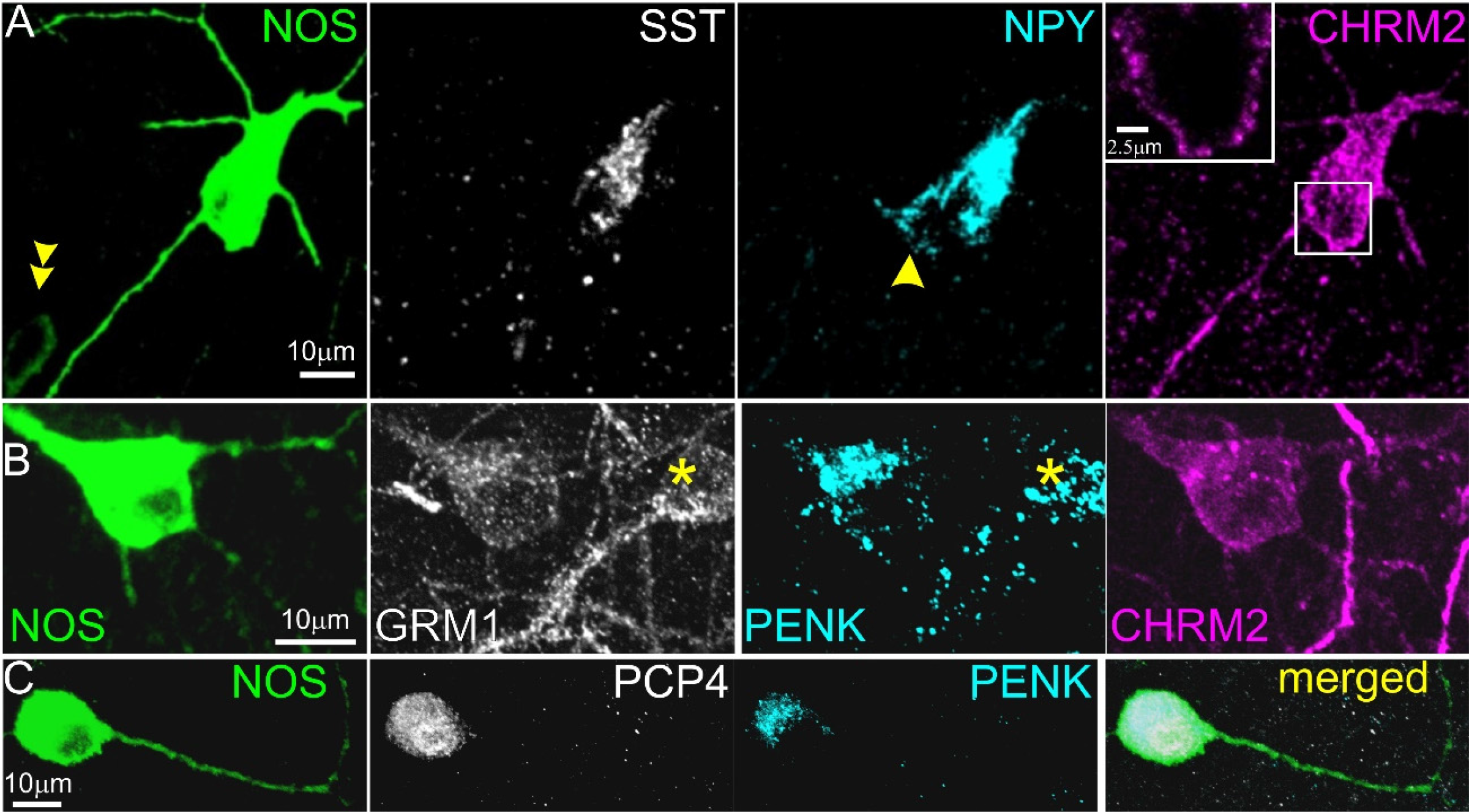
Immunohistochemical characterization of intensely NOS1-positive neurons. **A.** A large multipolar neuron in stratum pyramidale is strongly SST and NPY positive in the somatic Golgi apparatus and weakly positive for CHRM2 in the somato-dendritic plasma membrane (maximum intensity projection, z stack, height 11 µm; inset, maximum intensity projection of 3 optical slices, z stack height 2 µm). A smaller more weakly NOS1-positive cell (double arrow) in lower left is immunonegative for the other molecules; a second NPY positive cell (arrow) adjoining the NOS1+ neuron is immunonegative for the other three molecules. **B**. A NOS1-positive cell and another NOS1-immunonegative cell (asterisk) at the border of strartum radiatum and lacunosum-moleculare are both positive for GRM1 in the plasma membrane and PENK in the Golgi apparatus and in granules, but only the NOS1+ cell is immunopositive for CHRM2 (maximum intensity projection, z stack, height 10 µm). **C**. An intensely NOS1-positive cell in stratum radiatum is also positive for PCP4 in the cytoplasm and nucleus, and for PENK in the Golgi apparatus and in granules (maximum intensity projection, z stack, height 15 µm).

A second prediction of our classification was the expression of *Npy* in multiple subclasses of *Cck* cell, most notably the *Slc17a8* and *Calb1* expressing clusters of continent 8. This was unexpected, as NPY (at least at the protein level) has instead been traditionally associated with SST-expressing neurons and ivy/neurogliaform cells (Fuentealba et al., 2008a, Katona et al., 2014). Nevertheless, no studies to our knowledge have yet examined immunohistochemically whether the neuropeptides NPY and CCK can be colocalised in the same interneurons. We therefore tested this by double immunohistochemistry in *sr* and *slm* (**Figure 8A**, n=3 mice). Consistent with our predictions, 119 out of 162 (74±6%) of the cells immunopositive for pro-CCK were also positive for NPY (an additional 73 cells were positive for NPY only, which according to our identifications should represent neurogliaform and radiatum-retrohippocampal cells). A subset (176 cells) of NPY and/or pro-CCK immunopositive neurons were further tested for CALB1 in triple immunoreactions. As expected, nearly all CALB1-positive neurons were pro-CCK-positive (89±2%), and CALB1 immunoreactivity was seen in a subset of the cells containing both pro-CCK and NPY (27±3%). Additional triple immunohistochemistry for NPY, pro-CCK and SLC17A8

**Figure 8.**
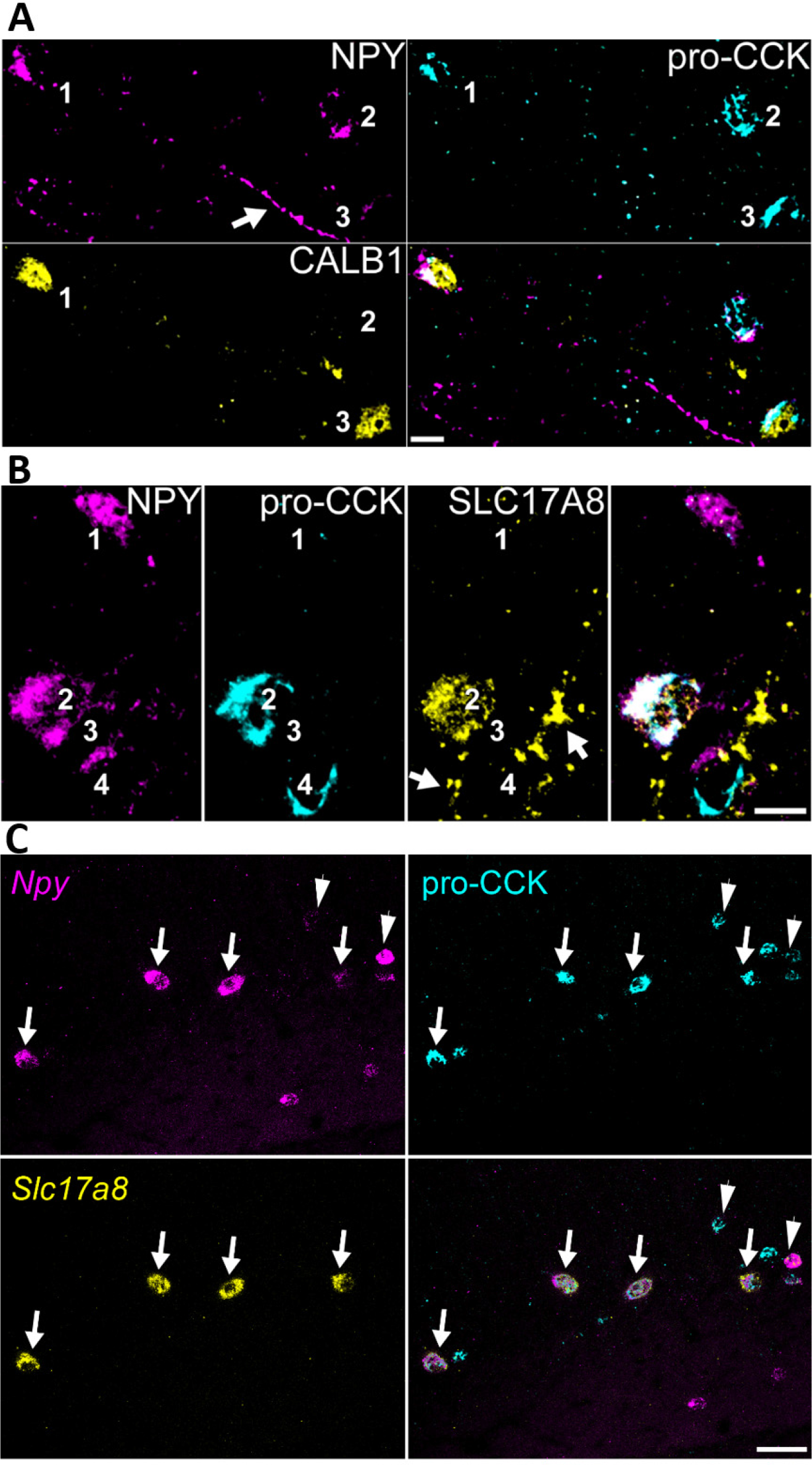
Confirmation of predicted co-localisation of NPY and pro-CCK. A, Interneurons at the sr/slm border immunopositive for both NPY and pro-CCK (cells 1 and 2), one of which (cell 1) is also immunopositive for CALB1. A third neuron is positive only for pro-CCK and CALB1 (cell 3). B, Interneurons at the sr/slm border immunopositive for NPY (cells 1-3), pro-CCK (cells 2 and 4) and SLC17A8 (VGLUT3, cell 2). Note SLC17A8-positive terminals targeting unlabelled cells (arrows). (a,b), Both NPY and pro-CCK are detected in the Golgi apparatus and endoplasmic reticulum surrounding cell nuclei, in addition, some axons are also immunopositive for NPY (see arrow in (a); average intensity projections, z stacks, height 6.3 µm and 10.4 µm, respectively). C, Combined double in situ hybridization and immunohistochemistry shows that nearly all Slc17a8-expressing cells also express Npy and are immunopositive for pro-CCK (arrows), but some Npy/pro-CCK cells do not express Slc17a8 (arrowheads). Scale bars: 10 µm (a,b), 50 µm (c).

(VGLUT3) revealed triple positive cells in *sr* and particularly at the *sr/slm* border, as predicted by the class *Cck.Cxcl14.Slc17a8* (**Figure 8B**). Due to the low level of somatic immunoreactivity for SLC17A8 (which as a vesicular transporter is primarily trafficked to axon terminals), we could not count these cells reliably; however of the cells that were unambiguously immunopositive for SLC17A8, in a majority we detected NPY. Additional analysis combining double *in situ* hybridization for *Slc17a8* and *Npy* with immunohistochemistry for pro-CCK (**Figure 8C**, n=3 mice) confirmed that the great majority of *Slc17a8*-expressing cells were also positive for *Npy* and pro-CCK (84±3%). As predicted by our identifications, the converse was not true: a substantial population of *Npy*/pro-CCK double-positive cells (57±7% of the total) did not show detectable *Slc17a8*, which we identify with dendrite-targeting neurons in the east of continent 8.

Several cell types in our classification expressed *Cxcl14*, a gene whose expression pattern in the Allen Brain Atlas shows localization largely at the *sr/slm* border. The *Cxcl14-*positive population includes all clusters of continent 8, which express *Cck* and contain subclusters expressing *Npy, Calb1, Reln*, and *Vip;* a subtype of CGE-derived neurogliaform cell that expresses *Reln* and *Npy* but lacks *Nos1* and expresses *Kit* at most weakly; as well as IS-1, IS-2, and radiatum-retrohippocampal cells. However, as all C*xcl14*-positive clusters lacked *Lhx6* we conclude they should be distinct from all MGE-derived neurons, including MGE-derived neurogliaform cells.

To test these predictions, we performed *in situ* hybridization for *Cxcl14* simultaneously with *in situ* hybridization or immunohistochemistry to detect *Reln, Npy*, CALB1, CCK, PVALB, *Sst, Nos1* and *Kit* (n=3 mice; **Figure 9**). In addition, we combined fluorescent *in situ* hybridization for *Cxcl14* with immunohistochemistry for YFP in *Lhx6-Cre/R26R-YFP* mice, which allows identification of developmental origin by marking MGE-derived interneurons (Fogarty et al., 2007). The results of these experiments were consistent with our hypotheses. We found that within CA1, *Cxcl14-*expressing cells were primarily located at the *sr/slm* border (71±3%), although a subpopulation of cells were also found in other layers. We found no overlap of *Cxcl14* with YFP in the *Lhx6-Cre/R26R-YFP* mouse, confirming the CGE origin of *Cxcl14* expressing neurons (**Figure 9A**); consistent with this finding, no overlap was seen with *Cxcl14* and *Sst* or *Pvalb* (data not shown). The majority of *Cxcl14*-positive cells expressed *Reln* (72±4%), accounting for 42±9% of Reln-expressing neurons (Substantial populations of *Reln*+/*Cxcl14-*cells located in *so* and *slm* likely represent O-LM and MGE-neurogliaform cells, respectively (**Figure 9B**). Indeed, although less than half of *Reln* cells were located at the R-LM border (44±1%), the great majority of *Reln+/Cxcl14+* cells were found there (88±6%). Consistent with the expected properties of continent 8 cells, a large fraction of the *Cxcl14* population were immunoreactive for pro-CCK (62±6%; **Figure 9C**), while substantial minorities were positive for CALB1 (29±2%; **Figure 9D**) or *Npy* (25±5%; **Figure 9E**). However, as expected from the lack of *Cxcl14* in MGE-derived neurogliaform and IS-3 cells, we observed no overlap of *Cxcl14* with *Nos1* (0 out of 209 cells; **Figure 9F)**; and very weak overlap with *Kit*, which is primarily expressed in clusters *Cacna2d1.Ndnf.Npy* and *Cacna2d1.Ndnf.Rgs10*, associated with the *Cxcl14-*negative CGE-neurogliaform population (1 of 264 cells respectively, from all mice; **Figure 9G**).

**Figure 9.**
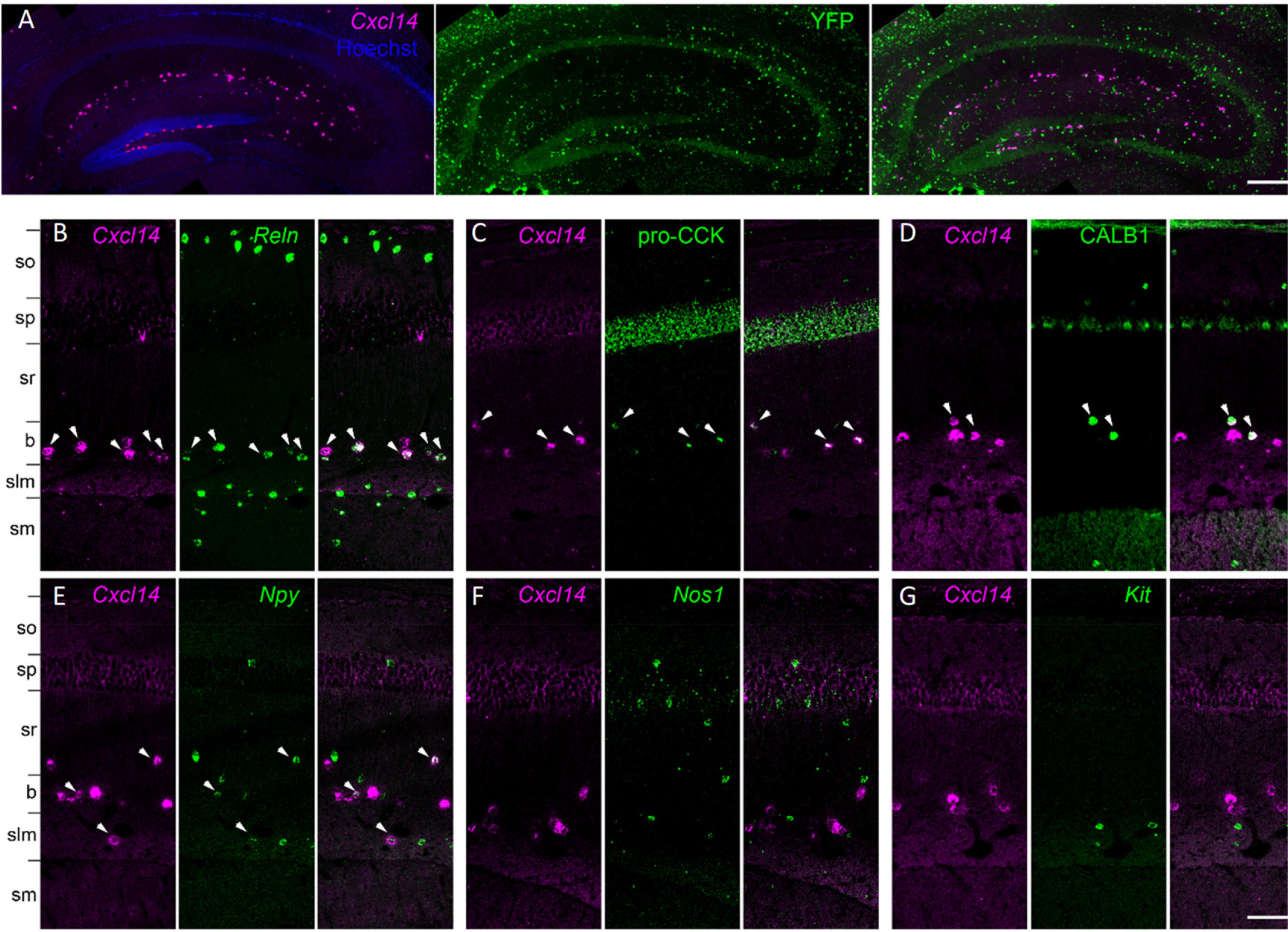
Analysis of *Cxcl14* co-expression patterns confirms predicted properties *Cck.Cxcl14* cells. **A**, *Cxcl14-*expressing cells are CGE-derived: in situ hybridization for *Cxcl14* combined with immunohistochemistry for YFP in the *Lhx6-Cre/R26R-YFP* mouse yields no double labelling. **B**, Double *in situ* hybridization for *Cxcl14* and *Reln* marks a population of neurons located primarily at the *sr/slm* border. Note *Reln* expression without *Cxcl14* in *so* and *slm*, likely reflecting O-LM and neurogliaform cells. **C-E**, Subsets of the *Cxcl14-*positive neurons are positive for pro-CCK or CALB1 (*in situ* hybridization plus immunohistochemistry), or *Npy* (double *in situ* hybridization). **(f, g)** No overlap was seen of *Cxcl14* with *Nos1* or *Kit*. In all panels, arrowheads indicate double-expressing neurons. Layer abbreviations: so, *stratum oriens;* sp, *stratum pyramidale;* sr, *stratum radiatum;* b, *sr/slm* border region; slm, *stratum lacunosum-moleculare;* sm, *stratum moleculare* of the dentate gyrus. Scale bars: 200 µm (a), 100 µm (b-g).

The cluster *Cck.Cxcl14.Vip* presented a puzzle, since *Cxcl14* is located primarily at the *sr/slm* border, whereas immunohistochemistry in rat has localized CCK/VIP basket cells to *sp* (Acsady et al., 1996a). Because *Cxcl14* expression can sometimes also be found in *sp*, we tested whether this cluster reflects *sp* cells, by combining in situ hybridization for *Cxcl14* with immunohistochemistry against VIP in mouse CA1 (**Figure 10)**. This revealed frequent co-expression at the *sr/slm* border (8 ±1% *Cxcl14* cells positive for *Vip*; 23 ±1% *Vip* cells positive for *Cxcl14*), but very few *Cxcl14* cells in *sp*, and essentially no double labelling (1 of 147 *Vip* cells in *sp* was weakly labelled for *Cxcl14)*. We therefore conclude that this cluster indeed represents a novel cell type located at the *sr/slm* border, expressing *Cck, Vip*, and *Cxcl14*.

**Figure 10.**
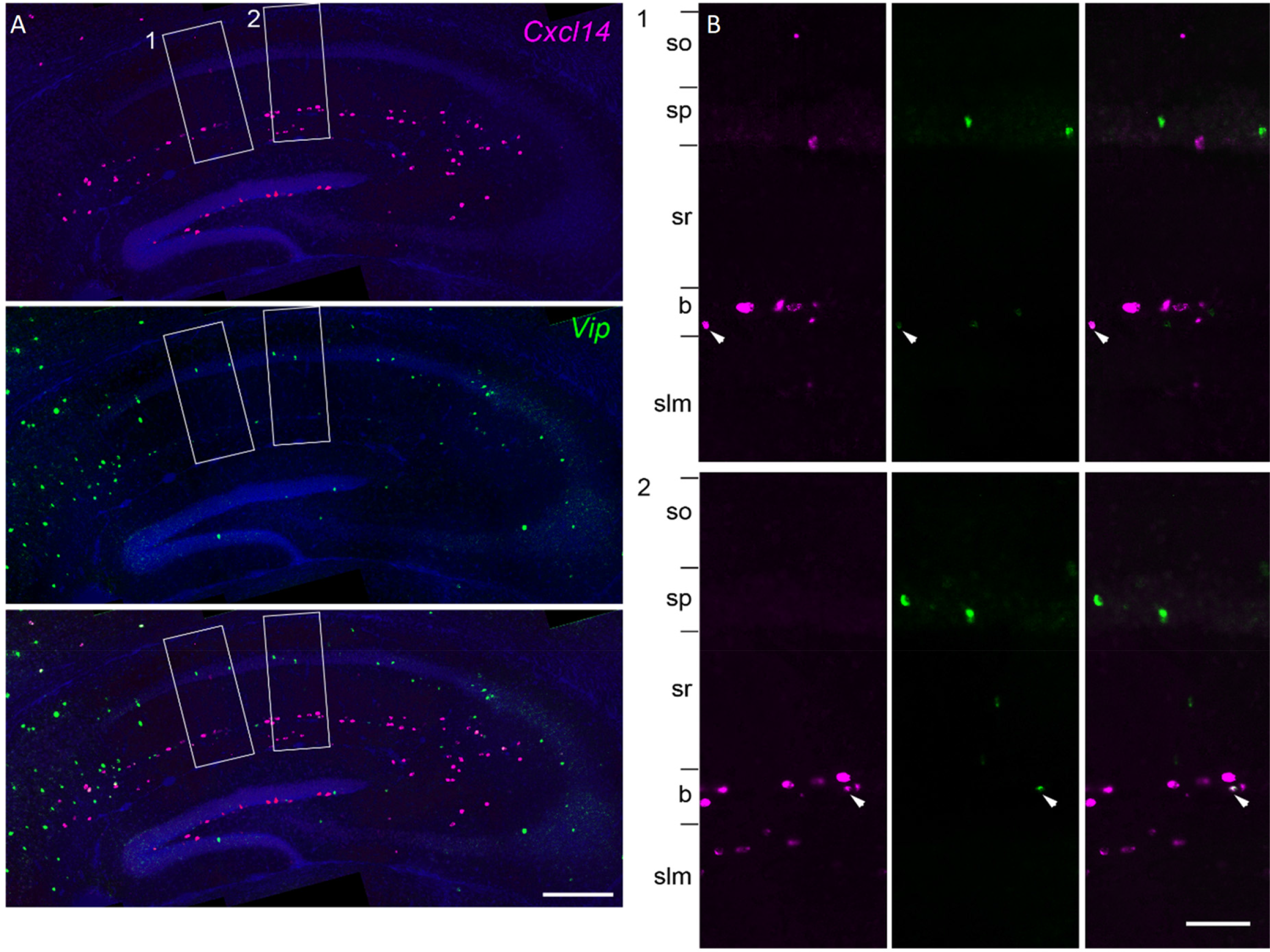
Overlap of *Cxcl14* and *Vip*. The class *Cck.Cxcl14.Vip* represented a puzzle: *Vip/Cck* cells had previously been reported in *sp*, but *Cxcl14* is detected primarily at the *sr/slm* border, although exceptional cells can be detected in *sp* also. **A**, double fluorescent *in situ* hybridization images reveal that the vast majority of cells co-expressing *Cxcl14* and *Vip* were found at the *sr/slm* border, confirming the location of this novel class. **B**, zoom into rectangles 1 and 2. Arrowheads: double-expressing cells.

## Discussion

The molecular architecture of CA1 interneurons has been intensively studied over the last decades, leading to the identification of 23 inhibitory classes. Our transcriptomic data showed a remarkable correspondence to this previous work, with all previously-described classes identified in our database. Our analysis also revealed a continuous mode of variability common across multiple cell types, eight hypothesized novel classes, as well as additional molecular subdivisions of previously described cell types.

Surprisingly, these data suggest that three previously described CA1 cell groups in fact represent a single cell class, a fact previously overlooked due to the limited combinations of molecules tested in prior work. The *Sst.Nos1* class is strongly positive for *Nos1*, and also expresses *Sst, Npy, Chrm2, Pcp4* and *Penk*, but unlike *Penk*-positive interneuron-selective cells of continent 9 lacks *Vip*. This class is homologous to the Int1 and Sst Chodl classes defined in isocortex, which have been identified with long-range projecting sleep-active neurons (Gerashchenko et al., 2008; Magno et al., 2012; Tasic et al., 2016; Zeisel et al., 2015). The three previously described classes identified with *Sst.Nos1* are: PENK-immunopositive neurons with projections to subiculum, that were shown to be VIP-negative, but not tested for SST or NOS1 (Fuentealba et al., 2008b); the NADPH diaphorase-labelled (i.e. strongly NOS1 positive) axons reported by Sik et al (1994) as projecting to CA3 and dentate, but not tested for SST or PENK; and the SST/NOS1 cells identified by Jinno and Kosaka (2004) in mouse, which were not tested for long-range projections or for PENK. While it remains possible that a larger transcriptomic sample of these rare neurons would reveal subclasses, our present data suggest that *Sst.Nos1* cells are a homogeneous population: the nbtSNE algorithm, BIC criterion, and further manual exploration failed to reveal any finer distinctions. We therefore suggest that they constitute a class of inhibitory neurons with diverse long-range projection targets. Interestingly, the targets of PENK-positive projection cells are most commonly PVALB-positive interneurons, unlike conventional IS cells, which preferentially target SST cells (Fuentealba et al., 2008a). As these cells are identified as sleep-active, this fact may provide an important clue to the mechanisms underlying sleep in cortical circuits.

The match between our transcriptomic analysis and previous immunohistochemical work (primarily in rat) is so close that it is simpler to describe the few areas of disagreement than the many areas of agreement. First, ACTN2 has been used as a neurogliaform marker in rat (Price et al., 2005), but was almost completely absent from any cell type of our database. We suggest this reflects a species difference, as previous attempts with multiple ACTN2 antibodies have been unsuccessful in mouse (JH-L, unpublished observations), and *Actn2* labelling is not detectable in the Allen atlas (Lein et al., 2007). Second, we observed *Calb2* in a subset of putative O-LM cells; these *Calb2-*expressing neurons typically also expressed *Calb1*. Such O-LM cells have not been described in rat (Katona et al., 2014), but CALB2/SST neurons have been observed in mouse isocortex (Tasic et al., 2016; Xu et al., 2006). A third inconsistency regards NCALD, which in rat was reported not to overlap with PVALB, SST, or NPY (Martínez-Guijarro et al., 1998), but did so in our data. Finally, it has previously been reported that a subset of O-LM cells show *Htr3a* expression (Chittajallu et al., 2013). In our data, we observed at most weak expression of *Htr3a* in *Sst* cells, and the cells showing it belonged to clusters identified as hippocamposeptal rather than O-LM cells.

Our analysis revealed several rare and presumably novel cell groups, although we cannot exclude that some of these were inadvertently included from neighboring areas such as subiculum (**Figure S10**). *Sst.Npy.Serpine2* and *Sst.Npy.Mgat4c*, which simultaneously expressed *Sst, Npy*, and *Reln* fit the expected expression pattern of neither O-LM nor hippocamposeptal cells; *Sst.Erbb4.Rgs10* is a distinct group related to *Pvalb* basket and bistratified cells; *Cck.Lypd1* formed a rare and highly distinct class expressing *Cck*, *Slc17a8*, and *Calb1*; *Ntng1.Synpr* showed an expression pattern with features of both *sr/slm Cck* neurons and projection cells; and *Cck.Cxcl14.Vip* represents a cell class strongly positive for both *Cck* and *Vip* located at the *sr/slm* border that appears to represent a pyramidal-rather than interneuron-targeting class. The analysis also revealed subdivisions of known types, such as the division of IS-3 cells into *Nos1* positive and negative groups, and the division of CGE-NGF cells into *Car4-* and *Cxcl14-*expressing subtypes. Finally, our data suggested that with more cells or deeper sequencing, even rarer types are likely to be found, as subsetting analysis showed a linear increase in the number of clusters with cell count and read depth, with little sign of saturation as yet. The data appeared to contain several novel cell types not containing enough cells to overcome the algorithm’s parsimony penalty, such as a small group of cells with features of both basket and axo-axonic cells located off the coast of continent 3; such cells have indeed been rarely encountered by quantitative electron microscopic analysis of synaptic targets in the rat (P. S., unpublished observations).

Latent factor analysis revealed a common continuum of gene expression across the database, suggesting a large “module” of genes that are co-regulated in multiple types of hippocampal interneuron. The latent factor differed between clusters, and clusters with larger latent factor values were identified with interneuron types targeting pyramidal cell somas or proximal dendrites (such as *Pvalb* or *Cck/Slc17a8* expressing basket cells), while those with low mean values were identified with interneurons targeting pyramidal distal dendrites (such as *Sst* or *Cck/Calb1* expressing dendrite-targeting cells) or other interneurons. Subtler differences in latent factor were found within clusters, suggesting that a similar continuum exists within cells of a single type. Genes positively correlated with the latent factor are associated with fast-spiking phenotype, presynaptic function, GABA release, and metabolism. Consistent with this expression pattern, perisomatic inhibitory cells show fast-spiking phenotypes and deliver powerful, accurately-timed inhibition (Hu et al., 2014), but interneurons targeting distal dendrites show slower-spiking patterns; presumably because distal inputs are subject to passive dendritic filtering, their presynaptic vesicle release does not need to be so accurately timed. I-S cells had the lowest mean values of the latent factor, consistent with their small axonal trees and metabolic machinery (Gulyás et al., 2006). The stronger expression of many neuropeptides in cells of low latent factor suggests that these slower, distal-targeting interneurons may also rely more heavily on neuropeptide signaling, for which slow firing rates support outputs transduced by slower G-protein coupled receptors. Interestingly, a study conducted independently of the present work identified enriched expression of a gene module similar to our latent factor in isocortical *Pvalb* neurons (Paul et al., 2017), and suggested it is controlled by the transcription factor PGC-1α (Lucas et al., 2010, 2014). Our results suggest that *Cck-*expressing basket cells have a similar expression pattern, and that more generally, expression of this module correlates with a neuron’s axonal target location.

Several novel genes correlating with the factor appear interesting candidates for future research, such as *Trp53i11, Yjefn3* and *Rgs10*, associated with faster spiking *Cck* cells; *Zcchc12* and *6330403K07Rik*, both associated with slower-firing cells of all classes; and *Fxyd6*, associated with slow-spiking, which may modulate ion exchange. Intriguingly, genes for neurofilaments and other intermediate filaments were positively correlated with the latent factor, while genes involved in actin processing were negatively correlated; we hypothesize that this might reflect a different cytoskeletal organization required for somatic- and dendritic-targeting neurons.

The question of how many cell classes a given neural circuit contains is often asked of transcriptomic analyses, but we argue this question will not have a clearly defined answer. For example, our data indicate no sharp dividing line between ivy cells and MGE-derived neurogliaform cells. Yet cells at the two ends of the continuum are clearly different: not only do their gene expression patterns differ substantially, but their different axonal targets indicate different roles in circuit function (Fuentealba et al., 2008b). In statistics, multiple criteria can be used to define how many clusters should be assigned to a dataset; a common approach (which is used by the ProMMT algorithm) is to consider a cluster indivisible if within-cluster fluctuations cannot be distinguished from random noise. Using this criterion, the number of clusters of CA1 interneurons increased with the number of cells and read depth analyzed, showing no sign of saturation in the current dataset. Furthermore, we observed several apparent rare classes that were too small to be assigned their own clusters at present, together with further subtle gradations within currently assigned clusters. The fact that we observed more clusters in CA1 than the 23 previously identified in isocortex (Tasic et al., 2016) should therefore not be taken as implying that CA1 is a more complex circuit, simply that our larger sample size and different clustering algorithm were able to detect finer distinctions. Indeed, our data suggest that while the divisions between the 10 major “continents” are unambiguous, the organization of gene expression within these continents is complex and subtle, and likely far more detailed than characterized by our present 49 clusters. An understanding this multiscale variability in gene expression in CA1 interneurons will be a key tool to understand the function of this circuit.

## Methods

### Single-cell RNA sequencing

#### Animals

*Slc32a1* (vesicular GABA transporter)-*Cre* BAC transgenic mice (Ogiwara et al., 2013) were crossed with a tdTomato reporter line to generate mice with fluorescently labelled inhibitory neurons. Both the *Slc32a1-Cre* and tdTomato mouse lines were of mixed B6 and CD1 backgrounds. Three of these mice were used for both the p28 and p63 cohorts; both males and females were used at each age. All experimental procedures followed the guidelines and recommendations of Swedish animal protection legislation and were approved by the local ethical committee for experiments on laboratory animals (Stockholms Norra Djurförsöksetiska nämnd, Sweden). The data are available on GEO under accession number GSE99888.

#### Single cell suspension and FACS

Dissection and single cell dissociation were carried out as described before (Marques et al. 2016), with slight alterations for P63 animals, where NMDG-HEPES based solution was used in all steps to enable better recovery of the aged cells (Tanaka, 2008). The NMDG-HEPES based cutting solution contained 93mM NMDG, 2.5mM KCl, 1.2mM NaH_2_PO_4_, 30mM NaHCO_3_, 20mM HEPES, 25mM Glucose, 5mM sodium ascorbate, 2mM thiourea, 3mM sodium pyruvate, 10mM MgSO_4_*7H_2_O, 0.5mM CACl_2_*2H_2_O and 12mM N-acetyl-L-cysteine; it was adjusted to pH 7.4 with 10N HCl. Mice were sacrificed by an overdose of Isoflurane and Ketamine/Xylazine, followed by transcardial perfusion through the left ventricle with artificial cerebrospinal fluid (aCSF) equilibrated in 95%O2 5%CO2 before use. The brain was removed and CA1 was microdissected from 300μm vibratome sections. Single-cell suspensions were prepared using Papain (Worthington) with 30min enzymatic digestion, followed by manual trituration with fire-polished Pasteur pipettes. The albumin density gradient was only performed for the p63 samples. On a BD FACSAria II, tdTomato-positive cells were sorted into oxygenated aCSF at 4°C, concentrated, inspected for viability, and counted.

To assess the accuracy of our dissection, we studied the gene expression patterns of simultaneously collected pyramidal cells using previously published genetic criteria (Figure **S8**). No cells exhibited expression patterns consistent with CA2 or CA3 (Cembrowski et al., 2016a), but a fraction of these cells (62 of 357 total excitatory neurons) expressed genes seen in a region stretching from the dorsomedial lip of CA1 to the subiculum. Although this result is consistent with dissection of only CA1 interneurons, we also cannot rule out the presence of a small number atypical interneuron classes located at the dorsomedial lip, or of inclusion of some subicular interneurons.

#### 10X Chromium mRNA-seq

Sorted suspensions were added to 10X Chromium RT mix aiming at 2500 cells recovered per experiment. Downstream cDNA synthesis (14 PCR cycles) and library preparation were carried out as instructed by the manufacturer (10X Genomics Chromium Single Cell Kit Version 1). Libraries were sequenced on the Illumina HiSeq2500 to an average depth 112 000 reads per cell (raw), yielding on average 3600 distinct molecules and 1700 genes per cell. Demultiplexed samples were aligned to the reference genome and converted to mRNA molecule counts using the “cellranger” pipeline version 1.1, provided by the manufacturer.

#### Normalization

Prior to many analyses (including clustering, latent factor analysis and nbtSNE) the expression vectors for each cell were normalized, so that each cell’s total RNA expression became equal to the total cellular RNA count averaged over all cells in the database. However, scatterplots of expression (Figure 6C) show unnormalized values.

#### Quality control

Cells showing abnormally high values of nuclear noncoding RNAs (*Meg3, Malat1, Snhg11*) or mitochondrial genes were discarded, as this can signify cell lysis. Cells were discarded if the summed normalized expression of these genes exceeded a threshold of 600.

### Immunohistochemistry (Oxford)

Six adult (20 weeks old) male C57BL/6J mice (Charles River, Oxford, UK) were perfusion fixed following anaesthesia and tissue preparation for immunofluorescence (Katona et al., 2014) and analysis using wide-field epifluorescence microscopy (Somogyi et al., 2004) were performed as described. The following primary antibodies were used: anti-calbindin (goat, Fronteir Inst, Af104); anti-pro-CCK (rabbit, 1:2000, Somogyi et al., 2004); anti-metabotropic glutamate receptor 1a (GRM1, rabbit, 1:1000; guinea pig, 1:500; gifts from Prof. M. Watanabe, Frontier Institute); anti-muscarinic acetylcholine receptor 2 (CHRM2, rat, 1:400, EMD Millipore Corporation, MAB367); anti-NOS1 (rabbit, 1:1000, EMD Millipore Corporation, AB5380; mouse, 1:1000, Sigma-Aldrich, N2280); anti-NPY (mouse, 1:5000, Abcam, #ab112473); anti-Purkinje cell protein 4 (PCP4, rabbit, 1:500, Santa Cruz Biotechnology, sc-74816); anti-pre-pro-enkephalin (PENK, guinea pig, 1:1000, gift from Takahiro Furuta, Kyoto University, Japan; rabbit, 1:5000, LifeSpan Biosciences, LS-C23084); anti-SST (sheep, 1:500, Fitzgerald Industries International, CR2056SP); anti-VGLUT3 (guinea pig, Somogyi et al 2004). Secondary antibodies were raised in donkey against immunoglobulin G of the species of origin of the primary antibodies and conjugated to Violet 421 (1:250); DyLight405 (1:250); Alexa 488 (1:1000); cyanine 3 (1:400); Alexa 647 (1:250); cyanine 5 (Cy5, 1:250). With the exception of donkey-antimouse-Alexa488 purchased from Invitrogen, all secondary antibodies were purchased from Stratech.

For cell counting, image stacks (212 × 212 μm area; 512 × 512 pixels; z stack height on average 12 μm) were acquired using LSM 710/AxioImager.Z1 (Carl Zeiss) laser scanning confocal microscope equipped with Plan-Apochromat 40x/1.3 Oil DIC M27 objective and controlled using ZEN (2008 v5.0 Black, Carl Zeiss). In a second set of sections, images were taken using Leitz DM RB (Leica) epifluorescence microscope equipped with PL Fluotar 40x/0.5 objective. Counting was performed either using ImageJ (v1.50b, Cell Counter plugin) on the confocal image stacks or OPENLAB software for the epifluorescence documentation. For the CCK counts, numbers were pooled from two separate reactions testing for a given combination of primary antibodies (n=3 mice each reaction, 2-3 sections each mouse) and reported as average values ± standard deviation. For the testing of intensely nNOS-positive neurons cells were selected using Leitz DM RB (Leica) epifluorescence microscope equipped with PL Fluotar 40x/0.5 objective. Cells were pooled from three separate reactions testing for a given combination of primary antibodies (n=3 mice each reaction, 2 sections each mouse) and reported as pooled data. Image processing was performed using ZEN (2012 Blue, Carl Zeiss), ImageJ (v1.51m, open source), Inkscape (0.92, open source) and Photoshop (CS5, Adobe).

### *In situ* hybridization (UCL)

Wild type (C57BL/6/CBA) male and female adult (P30) mice and *Lhx6-Cre^Tg^* transgenic mice were perfusion-fixed as previously described (Rubin et al., 2010), followed by immersion fixation overnight in 4% paraformaldehyde. Fixed samples were cryoprotected by overnight immersion in 20% sucrose, embedded in optimal cutting temperature (OCT) compound (Tissue Tek, Raymond Lamb Ltd Medical Supplies, Eastbourne, UK) and frozen on dry ice. 30 *μ*m cryosections were collected in DEPC-treated PBS and double *in situ* hybridization was carried out as described (Rubin et al., 2010). Probes used included either a *Cxcl14*-(digoxgenin)DIG RNA probe in combination with *Reln*-(fluorescein)FITC, *Npy*-FITC or *Sst*-FITC or *Vip*-FITC probes, or a *Cxcl14*-FITC probe with *Nos1*-DIG, *Kit*-DIG, *Scl17a8*-DIG, or *Pvalb*-DIG probes. DIG-labelled probes were detected with an anti-DIG-alkaline phosphatase (AP)-conjugated antibody followed by application of a Fast Red (Sigma) substrate. The first reaction was stopped by washing 3 × 10 min in PBS, and the sections were incubated with an anti-FITC-Peroxidase (POD)-conjugated antibody (1:1500 - Roche) overnight. The POD signal was developed by incubating the sections with Tyramide-FITC:amplification buffer (1:100, TSA™-Plus, Perkin Elmer) for 10 minutes, at room temperature. For immunohistochemistry after *in situ* hybridization the following antibodies were used: anti-Calbindin (rabbit, 1:1000, Swant, Bellinzona, Switzerland); anti-pro-CCK (rabbit, 1:2000, Somogyi et al., 2004); anti-GFP (chicken, 1:500, Aves Labs). All sections were counterstained with Hoechst 33258 dye (Sigma, 1000-fold dilution) and mounted with Dako Fluorescence Mounting Medium (DAKO).

For cell counts, images (at least two sections per mouse) were acquired on an epifluorescence miscroscope (Zeiss) with a 10x objective. Several images spanning the entire hippocampal CA1 were stitched using Microsoft Image Composite Editor. Cells were counted manually in the CA1 area including *sr* and *slm* and in a subregion spanning 100 *μ*m across the border between *sr* and *slm*, where most *Cxcl14*-positive cells are located. Confocal images (z stack height on average 25 μm, 2 *μ*m spacing) were taken on a Leica confocal microscope under a 10x objective and processed for contrast and brightness enhancement with Photoshop (CS5, Adobe). A final composite was generated in Adobe Illustrator (CS5, Adobe).

### Cluster analysis

#### Sparse Mixture Model

The ProMMT algorithm performs cluster analysis by modeling molecular counts by a mixture of sparse multivariate negative binomial distributions. Specifically, let x represent the *N_genes_*-dimensional vector summarizing the expression of all genes in a single cell. We model the probability distribution of x with a mixture model:

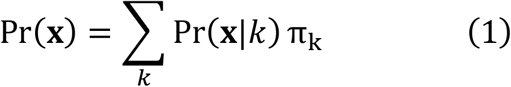

Here, *k* denotes a cell class, *p*(**x**|*k*) denotes the probability that a cell in this class will have expression vector **x**, and the “class prior” π_*k*_ represents the fraction of cells belonging to this class. To model *p*(**x**|*k*), we use the following distribution family:

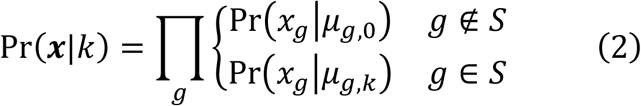

In this family, the distribution of all genes is modelled as conditionally independent within a class. The within-class distribution of each gene g depends on a single parameter µ_*g,k*_ (the mean level of the gene in that class). Furthermore, the distributions of only a subset *S* of genes are allowed to vary between classes, while the remainder are constrained to have a class-independent distribution with mean *µ*_*g*,O_. Taking *S* to have a fixed and small size *N_S_* ensures a “sparse model”, which can be fit robustly in high dimensions from only a small number of cells. Note that while the set S could in principle vary between classes, we have found that using a single set *S* for all classes provides good results.

#### Negative binomial distribution

To model the variability of each gene within a class, we use a negative binomial distribution. The negative binomial distribution is a model of count data with greater variance than the Poisson distribution, and is frequently used as a model for gene expression levels (Lu et al., 2005; Robinson and Smyth, 2008). The negative binomial is specified by two parameters, r and p, and has distribution

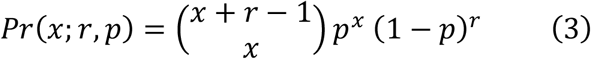

This distribution has mean 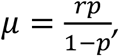, and for fixed *r* the maximum likelihood estimate of parameter *p* is 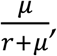 where *µ* is the sample mean. For fixed *r* the standard deviation of this distribution scales asymptotically linearly with its mean: 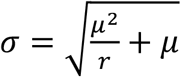. In contrast, the Poisson distribution has a smaller standard deviation, which scales with the square root of the mean.

We verified that a negative binomial with fixed *r* is appropriate for scRNA-seq data by considering a relatively homogeneous class (CA1 pyramidal cells; **Figure S2A**; data from Zeisel et al (2015)). This analysis confirmed that the negative binomial with *r* = 2 accurately modelled the relationship of standard deviation to mean in this data. The “wide” shape of the negative binomial distribution (**Figure S2B**) has a consequence that the absolute expression levels of a gene matters much less than whether the gene is expressed at all. Indeed, examining the symmetrized Kullback-Leibler divergence of negative binomials with different means (**Figure S2C**) – an indication of the penalty paid for misestimating the mean expression level – indicates that a much smaller penalty is paid for fitting a mean of 500 to a distribution whose actual mean is 1000, than to fitting a mean of 10 to a distribution whose actual mean is 0.

#### EM algorithm

To fit the model, we fix *r* = 2, and fit the parameters *S*, *µ*, and π by maximum likelihood. Because maximum likelihood fitting involves a sum over (unknown) class assignments, we use a standard Expectation-Maximization (EM) algorithm (Bishop, 2006; Dempster et al., 1977). We define z_*c,k*_ to be the expected value of an indicator variable taking the value 1 if cell c belongs to class k:

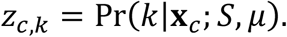

The algorithm alternates between an E step, where z_*c,k*_ is computed using the current values of the parameters *S* and *µ*, and an M step, where *S* and *µ* are optimized according to the current values of z_*c,k*_.

#### E-step

The E step is straightforward. Observe that

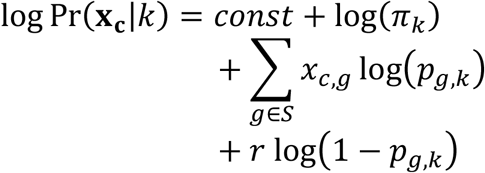

The constant term includes the contributions of all genes not in *S*, as well as the binomial coefficient from (3), none of which depend on the value of *k*, and therefore do not affect the result.

One can compute z_*c,k*_ from this using Bayes’ theorem; in practice, however we found that when the set *S* contains a reasonable number of genes (~100 or more) all values of z_*c,k*_ are close to 0 or 1, so there is little to lose by employing a much faster “hard EM” algorithm, in which for all cells c only a single winning *k_c_* has z_*c,k_c_*_ = 1, with all others 0.

#### M-step

In the M-step, we are given z_*c,k*_ and must find the set *S* of genes that are allowed to differ between classes, and their class means µ_*gk*_, by maximum likelihood. Although one might expect finding *S* to pose an intractable combinatorial optimization problem, it can in fact be solved quickly and exactly. The derivation below is for a hard EM algorithm; the soft case can be derived easily, but requires substantially more computation time, without a noticeable increase in performance.

We first define a quantity *L*_O_ to be the log likelihood of the data under a model where *S* = ∅, so all the expression of each genes *g* is determined by its grand mean *µ_g_*,_O_, independent of cluster assignments. Observe that

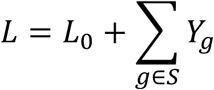

where

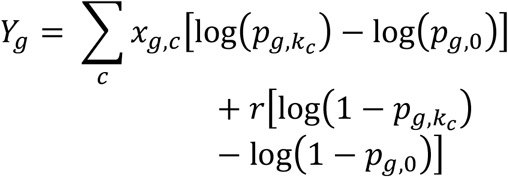

represents the gain in log likelihood obtained when the distribution of gene *g* is allowed to vary between classes. To compute the optimal value of the set *S*, we note that the values of *Y_g_* are independent of each other. Thus, the optimal set *S* is simply the *N_s_* genes with the largest values of *Y_g_*.

The maximum likelihood estimates of the negative binomial parameters *p_g,k_* are given by 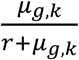, where *µ_g_*_,k_ denotes the average expression of gene *g* for the cells currently assigned to cluster *k*, and *µ_g_*_,O_ is the mean expression of gene *g* for all cells in the database. Because the negative binomial distribution can give zero likelihoods if any *µ_g,k_* = 0, we use a regularized mean estimate:

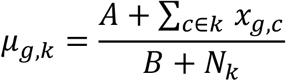

Where *N_k_* denotes the number of cells in cluster *k*, and the regularization parameters take the values *A* = 10^−4^, *B* = 1.

Finally, we compute the priors π_*k*_ as the fraction of cells *c* with *k_c_* = *k*, as is standard in EM.

#### BIC penalty

To automatically choose the number of clusters, we employed the BIC method (Schwarz, 1978), which for our model takes the form of a penalty 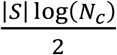 per cluster added to the log likelihood.

#### Cluster splitting

As is typical for cluster analysis, the likelihood function has multiple local maxima, and steps must be taken to ensure the algorithm does not become trapped in a suboptimal position. To do this, we use a heuristic that splits clusters that are poorly fit by a negative binomial distribution. The full clustering method consists of a divisive approach that alternates such splits with EM runs that then re-optimize the parameters.

For each cluster *k*, the splitting heuristic searches for genes *g* whose likelihood would be substantially increased if the cluster was split in two, according to whether the expression of gene *g* is above a threshold Θ_g_. Note that after splitting, the amount by which the log likelihood gain *Y_g_* changes can be written as

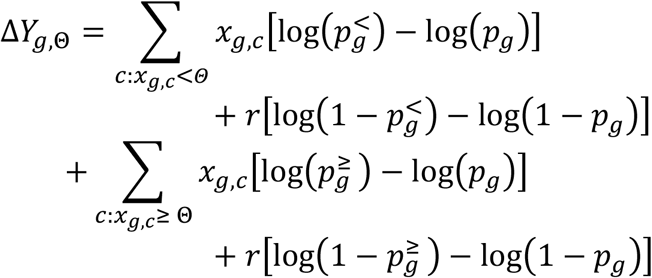

Here, *p_g_* represents the maximum-likelihood parameter for gene *g* in the cluster under consideration, 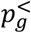 represents this parameter computed only for cells with *x_g_* < Θ_*g*_, and 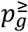 represents this parameter for cells with *x_g_* ≥ Θ_*g*_. The only values of Θ for which a split need be considered correspond to the expression levels of cells in the cluster, and ∆*Y_g_*_,0_ can therefore be rapidly computed for all *g* and Θ using cumulative summation, with computational cost linear in the size of the expression matrix.

#### Full algorithm

The full algorithm consists of repeatedly alternating the EM algorithm with cluster splitting and merging operations to escape from local maxima. The algorithm is initialized by assigning all cells to a single cluster.

On each iteration, all clusters are first split in two using the splitting heuristic. Specifically, for each cluster, ∆*Y_g_*_,Θ_ is computed for all *g* and Θ, and the optimal split points Θ_*g*_ are found for each gene. The ten genes giving top values of ∆*Y_g_*_,Θ_*g*__ are found. For each of them, the cluster is split, an EM algorithm run to convergence on the resulting cluster pair, and the split providing the highest increase in likelihood is kept. Once all clusters have been split, they are combined to produce a dataset with twice the original number of clusters, and the EM algorithm is run it, to allow points to be reassigned between the split clusters.

The iteration ends with a round of cluster pruning. For each cluster we compute the deletion loss: the decrease in log likelihood that would occur if all points in the cluster were reassigned to their second-best matching cluster. If this loss does not outweigh the BIC penalty, the cluster’s points are so reassigned, and EM is run on the full dataset. This process continues until no cluster’s deletion loss is smaller than the BIC penalty.

The algorithm is run for a set number of iterations (50 in the current case) and the final result corresponds to the clustering that gave highest score.

### Isolation metric

To measure how well separated each cluster is from its neighbors, we define an isolation metric equal to the deletion loss (described in the previous section), divided by *N_k_*log(2), where *N_k_* is the number of cells assigned to cluster *k*. This has an information-theoretic interpretation, as the number of additional bits that would be required to communicate the gene expression pattern of a cell in cluster *k*, using a code defined by the probability model if cluster *k* were deleted.

### Hierarchical cluster clustering

Each cluster produced by the EM algorithm is specified by a mean expression vector. To understand the relationship between these cluster means, we applied a clustering method to the clusters themselves. This was achieved using Ward’s method, with a distance matrix given by the K-L divergence between cluster means, weighted by the number of cells per cluster.

### nbtSNE algorithm

To visualize the locations of the cells we derived a variant of the tSNE algorithm (Maaten and Hinton, 2008) appropriate for data following a negative binomial distribution.

Stochastic neighbour embedding algorithms such as tSNE start by converting Euclidean distances between pairs of high-dimensional vectors *x_i_* into conditional probabilities according to a Gaussian distribution: 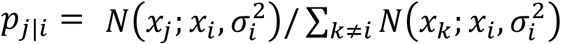 The tSNE algorithm then adjusts the locations of low-dimensional representation y_*i*_ in order to minimize the K-L divergence of a symmetrized p_*j*|*i*_, with a t-distribution on the y_*i*_.

The Gaussian distribution, however, is not the most appropriate choice for transcriptomic data. We found that we obtained better results using the same negative binomial distribution as in the ProMMT algorithm:

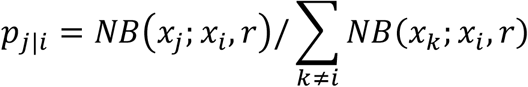

Where

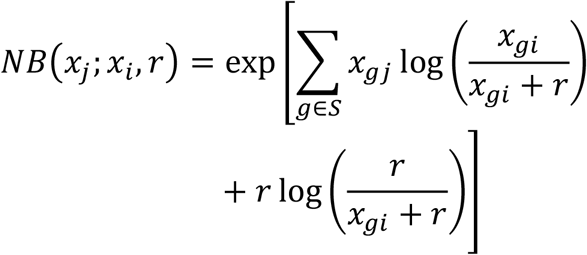

excluding a binomial coefficient that cancels when computing *p_j_*_|*i*_. The sum runs over the set of genes *g* that were chosen by the ProMMT algorithm.

In the original tSNE algorithm, variations in distance between the points x_i_ are overcome by adjusting the variance 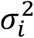 for each point *i* to achieve constant perplexity of the symmetrized conditional distributions. We took the same approach, finding a scale factor *λ_i_* for each cell *i* to ensure that the scaled symmetrized distribution

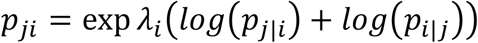

had a fixed perplexity of 15. This computation, and the implementation K-L minimization was achieved using Laurens van der Maaten’s drtoolbox (https://lvdmaaten.github.io/drtoolbox/). The algorithm was initialized by placing all points on a unit circle, with angular position determined by their parent cluster, linearly ordered by the hierarchical cluster clustering.

For comparison, we ran four other methods of tSNE analysis (**Figure S4)** using either all genes or the 150 genes found by ProMMT, and either a Euclidean metric or a Euclidean metric after log(1+x) transformation. Perplexity of 15 was again used and initialization was the same as before. Using all genes gave results that were difficult to interpret, particularly for log-transformed data, which we ascribe the noise arising from the large number of weakly expressed genes in the database. Using the gene subset provided more interpretable results, and combining the gene subset with log(1+x) transformation yielded results similar to nbtSNE, while Euclidean metric yielded less clear distinction of isolated classes such as *Cck.Lypd1* and *Sst.Nos1*. We conclude that the alignment of nbtSNE to the probability distribution of RNA counts of allows the algorithm to take into account differences between weakly expressed genes, and that a log(1+x) transformation approximates this probability distribution. We also conclude that gene subsetting prevents noise from the large number of genes that do not differ between classes from swamping the signal, and that this is particularly important with algorithms sensitive to changes in weakly expressed genes. We suggest that nbtSNE provides a principled probabilistic method for choosing the transformation and gene subset required for informative tSNE analysis.

### Negative binomial discriminant analysis

To investigate whether a pair of clusters were discretely separated or tiled a continuum we developed a method of cross-validated negative binomial discriminant analysis. This analysis assesses the separation of two clusters *k*_1_ and *k*_2_ by computing the log likelihood ratio for each cell to belong to the two clusters. It is simple to show that this ratio for a cell *c* is given by

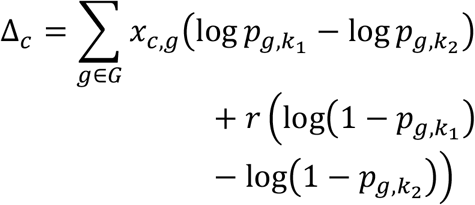

The sum runs over all genes g in the database, not just the set *S* found by the ProMMT algorithm.

The degree to which clusters *k*_1_ and *k*_2_ are discrete is visible by the bimodality of the histogram of ∆_*c*_, which can be quantified using a d’ statistic, 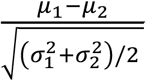, where *μ_i_* and *σ_i_* represent the mean and standard deviation of ∆_*c*_ for cells arising in cluster *i*. In this analysis, it is essential that the ratios ∆_*c*_ are computed on a separate “test set” of cells to the “training set” used to estimate *p_g,k_*, otherwise even a random division of a single homogeneous cluster would give an apparently bimodal histogram due to overfitting.

### Latent factor analysis

To model continuous variation between cells, we used a negative binomial latent factor model. The model is parametrized by two matrices, **W** and **F** of size *N_genes_* × *N_factors_* and *N_factors_* × *N_cells_*. The distribution of each cell follows a negative-binomial distribution with mean *µ_gc_* = *r* exp(∑*_f_ W_gf_ F_fc_*):

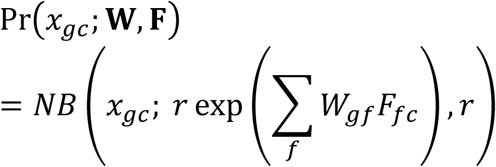

This corresponds to the natural parameterization of the negative binomial, *p* = 1/ (1 + exp(∑*_f_ W_gf_ F_fc_*)). As usual, we take a fixed value of *r* = 2. For the analysis described in this study, we use only a single latent factor, but add a second column to **F** of all ones to allow the mean expression level to vary between genes.

Given a dataset *x_gc_*, we fit the matrices **W** and **F** by maximum likelihood. As the negative binomial distribution with fixed *r* belongs to the exponential family, we can use the simple alternating method of Collins et al (2001). Note that we do not require a sparse algorithm because (unlike in clustering), the number of parameters is fixed. However to avoid instability, only genes that have reasonable expression levels in the database are kept (genes are included if at least 10 cells express at least 5 copies of the RNA), and a quadratic regularization penalty −50[|**W**|^2^ + |***F***|^2^] added to the log likelihood.

To relate the correlations of each gene with the latent factor to their gene ontology (GO) categories (supplementary table 2), we used the MGI mouse GO database (downloaded 2 April 2018), accessed via MATLAB’s bioinformatics toolbox. An enrichment score was computed for each GO term by summing the Spearman rank correlations of gene expression with the latent factor, over all genes annotated with that term.

## Author Contributions

KDH initiated collaboration (with JH-L), devised and implemented transcriptomic data analyses (Figs. 1-6, S1-S7), identified cell classes, wrote manuscript.

HH and NGS optimized protocols for adult mice and performed single cell sequencing.

LM performed FISH and IHC experiments and cell counts (Figs. 8,9 and S8C).

LK performed IHC experiments and cell counts (Figs. 7, S8A,B).

CBG bred animals, performed additional experiments, and analyzed data.

PS performed IHC experiments and cell counts (Figs. 7, S8A,B) contributed to cell class identification and manuscript writing.

NK supervised FIH and IHC experiments (Figs. 8,9 and S8C).

SL supervised single-cell sequencing.

JH-L initiated collaboration (with KDH), supervised single-cell experiments, contributed to manuscript writing.

## Acknowledgements

We thank T. Viney, A. Joshi, G. Unal, and B. Bekkouche for discussions and comments on the manuscript. This work was supported by the Wellcome Trust (108726 to KDH, NK, PS, SL, and JH-L), Medical Research Council (PS) European Research Council (261063 to SL), Swedish Research Council (STARGET to SL, and 2014-3863 to JH-L), StratNeuro (JH-L), Knut and Alice Wallenberg Foundation (SL) and European Union FP7/Marie Curie Actions (322304 to JH-L).

## REFERENCES

Acsady, L., Arabadzisz, D., and Freund, T.F. (1996a). Correlated morphological and neurochemical features identify different subsets of vasoactive intestinal polypeptide-immunoreactive interneurons in rat hippocampus. Neuroscience 73, 299–315.

Acsady, L., Gorcs, T.J., and Freund, T.F. (1996b). Different populations of vasoactive intestinal polypeptide-immunoreactive interneurons are specialized to control pyramidal cells or interneurons in the hippocampus. Neuroscience 73, 317–334.

Acsády, L., Pascual, M., Rocamora, N., Soriano, E., and Freund, T.F. (2000). Nerve growth factor but not neurotrophin-3 is synthesized by hippocampal GABAergic neurons that project to the medial septum. Neuroscience 98, 23–31.

Armstrong, C., and Soltesz, I. (2012). Basket cell dichotomy in microcircuit function: Basket cells as dichotomous microcircuit modulators. J. Physiol. 590, 683–694.

Armstrong, C., Krook-Magnuson, E., and Soltesz, I. (2012). Neurogliaform and Ivy Cells: A Major Family of nNOS Expressing GABAergic Neurons. Front. Neural Circuits 6.

Bezaire, M.J., and Soltesz, I. (2013). Quantitative assessment of CA1 local circuits: knowledge base for interneuron-pyramidal cell connectivity. Hippocampus 23, 751–785.

Bishop, C.M. (2006). Pattern Recognition and Machine Learning | Christopher Bishop | Springer (Springer verlag).

Blasco-Ibanez, J.M., Martinez-Guijarro, F.J., and Freund, T.F. (1998). Enkephalin-containing interneurons are specialized to innervate other interneurons in the hippocampal CA1 region of the rat and guinea-pig. Eur J Neurosci 10, 1784–1795.

Bouveyron, C., and Brunet-Saumard, C. (2014). Model-based clustering of high-dimensional data: A review. Comput. Stat. Data Anal. 71, 52–78.

Buhl, E.H., Halasy, K., and Somogyi, P. (1994). Diverse sources of hippocampal unitary inhibitory postsynaptic potentials and the number of synaptic release sites [see comments] [published erratum appears in Nature 1997 May 1;387(6628):106]. Nature 368, 823–828.

Cadwell, C.R., Palasantza, A., Jiang, X., Berens, P., Deng, Q., Yilmaz, M., Reimer, J., Shen, S., Bethge, M., Tolias, K.F., et al. (2016). Electrophysiological, transcriptomic and morphologic profiling of single neurons using Patch-seq. Nat. Biotechnol. 34, 199–203.

Cembrowski, M.S., Wang, L., Sugino, K., Shields, B.C., and Spruston, N. (2016a). Hipposeq: a comprehensive RNA-seq database of gene expression in hippocampal principal neurons. eLife 5, e14997.

Cembrowski, M.S., Bachman, J.L., Wang, L., Sugino, K., Shields, B.C., and Spruston, N. (2016b). Spatial Gene-Expression Gradients Underlie Prominent Heterogeneity of CA1 Pyramidal Neurons. Neuron 89, 351–368.

Chevée, M., Robertson, J.D., Cannon, G.H., Brown, S.P., and Goff, L.A. (2017). Variation in neuronal activity state, axonal projection target, and position principally define the transcriptional identity of individual neocortical projection neurons. bioRxiv 157149.

Chittajallu, R., Craig, M.T., McFarland, A., Yuan, X., Gerfen, S., Tricoire, L., Erkkila, B., Barron, S.C., Lopez, C.M., Liang, B.J., et al. (2013). Dual origins of functionally distinct O-LM interneurons revealed by differential 5-HT(3A)R expression. Nat Neurosci 16, 1598–1607.

Cho, J., Yu, N.-K., Choi, J.-H., Sim, S.-E., Kang, S.J., Kwak, C., Lee, S.-W., Kim, J., Choi, D.I., Kim, V.N., et al. (2015). Multiple repressive mechanisms in the hippocampus during memory formation. Science 350, 82–87.

Cohen, S.M., Ma, H., Kuchibhotla, K.V., Watson, B.O., Buzsáki, G., Froemke, R.C., and Tsien, R.W. (2016). Excitation-Transcription Coupling in Parvalbumin-Positive Interneurons Employs a Novel CaM Kinase-Dependent Pathway Distinct from Excitatory Neurons. Neuron 90, 292–307.

Collins, M., Dasgupta, S., and Schapire, R.E. (2001). A generalization of principal component analysis to the exponential family. In Advances in Neural Information Processing Systems, (MIT Press), p.

Cope, D.W., Maccaferri, G., Marton, L.F., Roberts, J.D., Cobden, P.M., and Somogyi, P. (2002). Cholecystokinin-immunopositive basket and Schaffer collateral-associated interneurones target different domains of pyramidal cells in the CA1 area of the rat hippocampus. Neuroscience 109, 63–80.

Daw, M.I., Tricoire, L., Erdelyi, F., Szabo, G., and McBain, C.J. (2009). Asynchronous transmitter release from cholecystokinin-containing inhibitory interneurons is widespread and target-cell independent. J Neurosci 29, 11112–11122.

Dehorter, N., Ciceri, G., Bartolini, G., Lim, L., del Pino, I., and Marin, O. (2015). Tuning of fast-spiking interneuron properties by an activity-dependent transcriptional switch. Science 349, 1216–1220.

Dempster, A.P., Laird, N.M., and Rubin, D.B. (1977). Maximum Likelihood from Incomplete Data via the EM Algorithm. J. R. Stat. Soc. Ser. B Methodol. 39, 1–38.

Donato, F., Rompani, S.B., and Caroni, P. (2013). Parvalbumin-expressing basket-cell network plasticity induced by experience regulates adult learning. Nature 504, 272–276.

Dudok, B., Barna, L., Ledri, M., Szabó, S.I., Szabadits, E., Pintér, B., Woodhams, S.G., Henstridge, C.M., Balla, G.Y., Nyilas, R., et al. (2015). Cell-specific STORM super-resolution imaging reveals nanoscale organization of cannabinoid signaling. Nat. Neurosci. 18, 75–86.

Ecker, J.R., Geschwind, D.H., Kriegstein, A.R., Ngai, J., Osten, P., Polioudakis, D., Regev, A., Sestan, N., Wickersham, I.R., and Zeng, H. (2017). The BRAIN Initiative Cell Census Consortium: Lessons Learned toward Generating a Comprehensive Brain Cell Atlas. Neuron 96, 542–557.

Ferraguti, F., Klausberger, T., Cobden, P., Baude, A., Roberts, J.D., Szucs, P., Kinoshita, A., Shigemoto, R., Somogyi, P., and Dalezios, Y. (2005). Metabotropic glutamate receptor 8-expressing nerve terminals target subsets of GABAergic neurons in the hippocampus. J Neurosci 25, 10520–10536.

Frazer, S., Prados, J., Niquille, M., Cadilhac, C., Markopoulos, F., Gomez, L., Tomasello, U., Telley, L., Holtmaat, A., Jabaudon, D., et al. (2017). Transcriptomic and anatomic parcellation of 5-HT3AR expressing cortical interneuron subtypes revealed by single-cell RNA sequencing. Nat. Commun. 8, 14219.

Freund, T.F., and Buzsaki, G. (1996). Interneurons of the hippocampus. Hippocampus 6, 347–470.

Fuentealba, P., Tomioka, R., Dalezios, Y., Marton, L.F., Studer, M., Rockland, K., Klausberger, T., and Somogyi, P. (2008a). Rhythmically active enkephalin-expressing GABAergic cells in the CA1 area of the hippocampus project to the subiculum and preferentially innervate interneurons. J Neurosci 28, 10017–10022.

Fuentealba, P., Begum, R., Capogna, M., Jinno, S., Marton, L.F., Csicsvari, J., Thomson, A., Somogyi, P., and Klausberger, T. (2008b). Ivy cells: a population of nitric-oxide-producing, slow-spiking GABAergic neurons and their involvement in hippocampal network activity. Neuron 57, 917–929.

Gerashchenko, D., Wisor, J.P., Burns, D., Reh, R.K., Shiromani, P.J., Sakurai, T., de la Iglesia, H.O., and Kilduff, T.S. (2008). Identification of a population of sleep-active cerebral cortex neurons. Proc. Natl. Acad. Sci. U. S. A. 105, 10227–10232.

Gulyás, A.I., and Freund, T.F. (1996). Pyramidal cell dendrites are the primary targets of calbindin D28k-immunoreactive interneurons in the hippocampus. Hippocampus 6, 525–534.

Gulyás, A.I., Hájos, N., and Freund, T.F. (1996). Interneurons containing calretinin are specialized to control other interneurons in the rat hippocampus. J. Neurosci. Off. J. Soc. Neurosci. 16, 3397–3411.

Gulyás, A.I., Buzsáki, G., Freund, T.F., and Hirase, H. (2006). Populations of hippocampal inhibitory neurons express different levels of cytochrome c. Eur. J. Neurosci. 23, 2581–2594.

Habib, N., Li, Y., Heidenreich, M., Swiech, L., Avraham-Davidi, I., Trombetta, J.J., Hession, C., Zhang, F., and Regev, A. (2016). Div-Seq: Single-nucleus RNA-Seq reveals dynamics of rare adult newborn neurons. Science 353, 925–928.

Habib, N., Basu, A., Avraham-Davidi, I., Burks, T., Choudhury, S.R., Aguet, F., Gelfand, E., Ardlie, K., Weitz, D.A., Rozenblatt-Rosen, O., et al. (2017). DroNc-Seq: Deciphering cell types in human archived brain tissues by massively-parallel single nucleus RNA-seq. bioRxiv.

Hefft, S., and Jonas, P. (2005). Asynchronous GABA release generates long-lasting inhibition at a hippocampal interneuron–principal neuron synapse. Nat. Neurosci. 8, 1319–1328.

Hu, H., Gan, J., and Jonas, P. (2014). Interneurons. Fast-spiking, parvalbumin+ GABAergic interneurons: from cellular design to microcircuit function. Science 345, 1255263.

Jiang, X., Wang, G., Lee, A.J., Stornetta, R.L., and Zhu, J.J. (2013). The organization of two new cortical interneuronal circuits. Nat Neurosci 16, 210–218.

Jiang, X., Shen, S., Cadwell, C.R., Berens, P., Sinz, F., Ecker, A.S., Patel, S., and Tolias, A.S. (2015). Principles of connectivity among morphologically defined cell types in adult neocortex. Science 350, aac9462.

Jinno, S. (2009). Structural organization of long-range GABAergic projection system of the hippocampus. Front Neuroanat 3, 13.

Jinno, S., and Kosaka, T. (2004). Patterns of colocalization of neuronal nitric oxide synthase and somatostatin-like immunoreactivity in the mouse hippocampus: quantitative analysis with optical disector. Neuroscience 124, 797–808.

Jinno, S., Klausberger, T., Marton, L.F., Dalezios, Y., Roberts, J.D., Fuentealba, P., Bushong, E.A., Henze, D., Buzsaki, G., and Somogyi, P. (2007). Neuronal diversity in GABAergic long-range projections from the hippocampus. J Neurosci 27, 8790–8804.

Katona, L., Lapray, D., Viney, T.J., Oulhaj, A., Borhegyi, Z., Micklem, B.R., Klausberger, T., and Somogyi, P. (2014). Sleep and movement differentiates actions of two types of somatostatin-expressing GABAergic interneuron in rat hippocampus. Neuron 82, 872–886.

Katona, L., Micklem, B., Borhegyi, Z., Swiejkowski, D.A., Valenti, O., Viney, T.J., Kotzadimitriou, D., Klausberger, T., and Somogyi, P. (2017). Behavior-dependent activity patterns of GABAergic long-range projecting neurons in the rat hippocampus. Hippocampus 27, 359–377.

Klausberger, T., and Somogyi, P. (2008). Neuronal diversity and temporal dynamics: the unity of hippocampal circuit operations. Science 321, 53–57.

Klausberger, T., Marton, L.F., O’Neill, J., Huck, J.H., Dalezios, Y., Fuentealba, P., Suen, W.Y., Papp, E., Kaneko, T., Watanabe, M., et al. (2005). Complementary roles of cholecystokinin- and parvalbumin-expressing GABAergic neurons in hippocampal network oscillations. J Neurosci 25, 9782–9793.

Lee, S.-H., Földy, C., and Soltesz, I. (2010). Distinct endocannabinoid control of GABA release at perisomatic and dendritic synapses in the hippocampus. J. Neurosci. Off. J. Soc. Neurosci. 30, 7993–8000.

Lein, E.S., Hawrylycz, M.J., Ao, N., Ayres, M., Bensinger, A., Bernard, A., Boe, A.F., Boguski, M.S., Brockway, K.S., Byrnes, E.J., et al. (2007). Genome-wide atlas of gene expression in the adult mouse brain. Nature 445, 168–176.

Losonczy, A., Zhang, L., Shigemoto, R., Somogyi, P., and Nusser, Z. (2002). Cell type dependence and variability in the short-term plasticity of EPSCs in identified mouse hippocampal interneurones. J. Physiol. 542, 193–210.

Lu, J., Tomfohr, J.K., and Kepler, T.B. (2005). Identifying differential expression in multiple SAGE libraries: an overdispersed log-linear model approach. BMC Bioinformatics 6, 165.

Lucas, E.K., Markwardt, S.J., Gupta, S., Meador-Woodruff, J.H., Lin, J.D., Overstreet-Wadiche, L., and Cowell, R.M. (2010). Parvalbumin Deficiency and GABAergic Dysfunction in Mice Lacking PGC-1α. J. Neurosci. 30, 7227–7235.

Lucas, E.K., Dougherty, S.E., McMeekin, L.J., Reid, C.S., Dobrunz, L.E., West, A.B., Hablitz, J.J., and Cowell, R.M. (2014). PGC-1α Provides a Transcriptional Framework for Synchronous Neurotransmitter Release from Parvalbumin-Positive Interneurons. J. Neurosci. 34, 14375–14387.

Maaten, L. van der, and Hinton, G. (2008). Visualizing Data using t-SNE. J. Mach. Learn. Res. 9, 2579–2605.

Macosko, E.Z., Basu, A., Satija, R., Nemesh, J., Shekhar, K., Goldman, M., Tirosh, I., Bialas, A.R., Kamitaki, N., Martersteck, E.M., et al. (2015). Highly Parallel Genome-wide Expression Profiling of Individual Cells Using Nanoliter Droplets. Cell 161, 1202–1214.

Magno, L., Oliveira, M.G., Mucha, M., Rubin, A.N., and Kessaris, N. (2012). Multiple embryonic origins of nitric oxide synthase-expressing GABAergic neurons of the neocortex. Front Neural Circuits 6, 65.

Mardinly, A.R., Spiegel, I., Patrizi, A., Centofante, E., Bazinet, J.E., Tzeng, C.P., Mandel-Brehm, C., Harmin, D.A., Adesnik, H., Fagiolini, M., et al. (2016). Sensory experience regulates cortical inhibition by inducing IGF1 in VIP neurons. Nature 531, 371–375.

Markram, H., Toledo-Rodriguez, M., Wang, Y., Gupta, A., Silberberg, G., and Wu, C. (2004). Interneurons of the neocortical inhibitory system. Nat.Rev.Neurosci. 5, 793–807.

Martínez-Guijarro, F.J., Briñón, J.G., Blasco-Ibáñez, J.M., Okazaki, K., Hidaka, H., and Alonso, J.R. (1998). Neurocalcin-immunoreactive cells in the rat hippocampus are GABAergic interneurons. Hippocampus 8, 2–23.

Miyashita, T., and Rockland, K.S. (2007). GABAergic projections from the hippocampus to the retrosplenial cortex in the rat. Eur J Neurosci 26, 1193–1204.

Ogiwara, I., Iwasato, T., Miyamoto, H., Iwata, R., Yamagata, T., Mazaki, E., Yanagawa, Y., Tamamaki, N., Hensch, T.K., Itohara, S., et al. (2013). Nav1.1 haploinsufficiency in excitatory neurons ameliorates seizure-associated sudden death in a mouse model of Dravet syndrome. Hum. Mol. Genet. 22, 4784–4804.

Parra, P., Gulyas, A.I., and Miles, R. (1998). How many subtypes of inhibitory cells in the hippocampus? Neuron 20, 983–993.

Paul, A., Crow, M., Raudales, R., Gillis, J., and Huang, Z.J. (2017). Transcriptional Architecture of Synaptic Communication Delineates Cortical GABAergic Neuron Identity. bioRxiv 180034.

Pawelzik, H., Hughes, D.I., and Thomson, A.M. (2002). Physiological and morphological diversity of immunocytochemically defined parvalbumin- and cholecystokinin-positive interneurones in CA1 of the adult rat hippocampus. J. Comp. Neurol. 443, 346–367.

Pelkey, K.A., Chittajallu, R., Craig, M.T., Tricoire, L., Wester, J.C., and McBain, C.J. (2017). Hippocampal GABAergic Inhibitory Interneurons. Physiol. Rev. 97, 1619–1747.

Price, C.J., Cauli, B., Kovacs, E.R., Kulik, A., Lambolez, B., Shigemoto, R., and Capogna, M. (2005). Neurogliaform neurons form a novel inhibitory network in the hippocampal CA1 area. J. Neurosci. Off. J. Soc. Neurosci. 25, 6775–6786.

Robinson, M.D., and Smyth, G.K. (2008). Small-sample estimation of negative binomial dispersion, with applications to SAGE data. Biostatistics 9, 321–332.

Schwarz, G. (1978). Estimating the Dimension of a Model. Ann. Stat. 6, 461–464.

Sik, A., Ylinen, A., Penttonen, M., and Buzsaki, G. (1994). Inhibitory CA1-CA3-hilar region feedback in the hippocampus. Science 265, 1722–1724.

Somogyi, P. (2010). Hippocampus: intrinsic organization. In Handbook of Brain Microcircuits, G.M. Shepherd, and S. Grillner, eds. (New York: Oxford University Press), p.

Somogyi, J., Baude, A., Omori, Y., Shimizu, H., El Mestikawy, S., Fukaya, M., Shigemoto, R., Watanabe, M., and Somogyi, P. (2004). GABAergic basket cells expressing cholecystokinin contain vesicular glutamate transporter type 3 (VGLUT3) in their synaptic terminals in hippocampus and isocortex of the rat. Eur J Neurosci 19, 552–569.

Spiegel, I., Mardinly, A.R., Gabel, H.W., Bazinet, J.E., Couch, C.H., Tzeng, C.P., Harmin, D.A., and Greenberg, M.E. (2014). Npas4 regulates excitatory-inhibitory balance within neural circuits through cell-type-specific gene programs. Cell 157, 1216–1229.

Takács, V.T., Freund, T.F., and Gulyás, A.I. (2008). Types and synaptic connections of hippocampal inhibitory neurons reciprocally connected with the medial septum. Eur. J. Neurosci. 28, 148–164.

Tasic, B., Menon, V., Nguyen, T.N., Kim, T.K., Jarsky, T., Yao, Z., Levi, B., Gray, L.T., Sorensen, S.A., Dolbeare, T., et al. (2016). Adult mouse cortical cell taxonomy revealed by single cell transcriptomics. Nat. Neurosci. 19, 335–346.

Tricoire, L., Pelkey, K.A., Daw, M.I., Sousa, V.H., Miyoshi, G., Jeffries, B., Cauli, B., Fishell, G., and McBain, C.J. (2010). Common origins of hippocampal Ivy and nitric oxide synthase expressing neurogliaform cells. J Neurosci 30, 2165–2176.

Tricoire, L., Pelkey, K.A., Erkkila, B.E., Jeffries, B.W., Yuan, X., and McBain, C.J. (2011). A blueprint for the spatiotemporal origins of mouse hippocampal interneuron diversity. J Neurosci 31, 10948–10970.

Tyan, L., Chamberland, S., Magnin, E., Camire, O., Francavilla, R., David, L.S., Deisseroth, K., and Topolnik, L. (2014). Dendritic inhibition provided by interneuron-specific cells controls the firing rate and timing of the hippocampal feedback inhibitory circuitry. J Neurosci 34, 4534–4547.

Usoskin, D., Furlan, A., Islam, S., Abdo, H., Lonnerberg, P., Lou, D., Hjerling-Leffler, J., Haeggstrom, J., Kharchenko, O., Kharchenko, P.V., et al. (2015). Unbiased classification of sensory neuron types by large-scale single-cell RNA sequencing. Nat Neurosci 18, 145–153.

Viney, T.J., Lasztoczi, B., Katona, L., Crump, M.G., Tukker, J.J., Klausberger, T., and Somogyi, P. (2013). Network state-dependent inhibition of identified hippocampal CA3 axo-axonic cells in vivo. Nat. Neurosci. 16, 1802–1811.

Vruwink, M., Schmidt, H.H., Weinberg, R.J., and Burette, A. (2001). Substance P and nitric oxide signaling in cerebral cortex: anatomical evidence for reciprocal signaling between two classes of interneurons. J. Comp. Neurol. 441, 288–301.

Wheeler, D.W., White, C.M., Rees, C.L., Komendantov, A.O., Hamilton, D.J., and Ascoli, G.A. (2015). Hippocampome.org: a knowledge base of neuron types in the rodent hippocampus. Elife 4.

Xu, X., Roby, K.D., and Callaway, E.M. (2006). Mouse cortical inhibitory neuron type that coexpresses somatostatin and calretinin. J. Comp. Neurol. 499, 144–160.

Zeisel, A., Munoz-Manchado, A.B., Codeluppi, S., Lonnerberg, P., La Manno, G., Jureus, A., Marques, S., Munguba, H., He, L., Betsholtz, C., et al. (2015). Brain structure. Cell types in the mouse cortex and hippocampus revealed by single-cell RNA-seq. Science 347, 1138–1142.

